# Differential functional reorganization of ventral and dorsal visual pathways following childhood hemispherectomy

**DOI:** 10.1101/2023.08.03.551494

**Authors:** Vladislav Ayzenberg, Michael C. Granovetter, Sophia Robert, Christina Patterson, Marlene Behrmann

## Abstract

Hemispherectomy is a surgical procedure in which an entire hemisphere of a patient’s brain is resected or functionally disconnected to manage seizures in individuals with drug-resistant epilepsy. Despite the extensive loss of input from both ventral and dorsal visual pathways of one hemisphere, pediatric patients who have undergone hemispherectomy show a remarkably high degree of perceptual function across many domains. In the current study, we sought to understand the extent to which functions of the ventral and dorsal visual pathways reorganize to the contralateral hemisphere following childhood hemispherectomy. To this end, we collected fMRI data from an equal number of left and right hemispherectomy patients who completed tasks that typically elicit lateralized responses from the ventral or the dorsal pathway, namely, word (left ventral), face (right ventral), tool (left dorsal), and global form (right dorsal) perception. Overall, there was greater evidence of functional reorganization in the ventral pathway than in the dorsal pathway. Importantly, because ventral and dorsal reorganization was tested in the very same patients, these results cannot be explained by idiosyncratic factors such as disease etiology, age at the time of surgery, or age at testing. These findings suggest that because the dorsal pathway may mature earlier, it may have a shorter developmental window of plasticity than the ventral pathway and, hence, be less malleable.

## Introduction

In adulthood, damage to higher-level regions of visual cortex, for example, following traumatic insult, stroke or surgery, causes profound visual processing deficits. The anatomical origins of many of these deficits are lateralized, such that damage to the left versus right hemisphere may cause a distinct pattern of impairment. For instance, in the ventral visual pathway, damage to portions of the left hemisphere of the ventral occipital temporal cortex (VOTC) commonly results in deficits in perceiving and recognizing written text (Behrmann, Black, & Bub, 1990; Behrmann, Plaut, & Nelson, 1998; Cohen et al., 2003), whereas damage to portions of the right hemisphere commonly leads to deficits in face recognition (Albonico & Barton, 2019; Landis, Cummings, Christen, Bogen, & Imhof, 1986). Similarly, in addition to impairing motor movements (Goodale et al., 1994; Goodale & Milner, 1992), lateralized damage to the dorsal pathway can cause perceptual deficits. For instance, damage to the posterior parietal cortex (PPC) of the left hemisphere may cause deficits in understanding how to use objects as tools (Garcea et al., 2018; Johnson-Frey, 2004), whereas damage to portions of the right hemisphere may cause deficits in perceiving global object form (Karakose-Akbiyik, Schubert, & Caramazza, 2023; Romei, Driver, Schyns, & Thut, 2011). In adults, these deficits may be long lasting and, even with intervention, may never recover fully (Behrmann & McLeod, 1995).

In contrast to adults, however, accumulating evidence suggests that similar (or even greater) disruptions to cortex in childhood may not result in deficits as severe as those observed in adulthood (Bourne, 2010; Granovetter, Patterson, & Behrmann, under revision; Kolb & Gibb, 2011). Indeed, children who undergo large-scale surgical resections of VOTC show strong recognition abilities and demonstrate neural reorganization of word and face representations to spared regions of cortex . For instance, pediatric patients who had portions of their left VOTC removed, encompassing regions crucial for reading (i.e., visual word form area; VWFA), nevertheless learn to read and evince selectivity to written words in their intact right hemisphere (Liu, Freud, Patterson, & Behrmann, 2019). Similarly, patients who had portions of their right VOTC removed, encompassing regions crucial for face perception (i.e., fusiform face area; FFA), nevertheless recognize faces and demonstrate face selectivity in their preserved left hemisphere (Liu & Behrmann, 2017). Indeed, pediatric patients with left or right ventral resections show surprisingly high accuracy on face and word recognition tasks that is only modestly, albeit statistically, lower than controls (Granovetter, Robert, Ettensohn, & Behrmann, 2022). Thus, unlike in adulthood, the developing ventral pathway may be sufficiently plastic to permit reorganization following large-scale damage.

Much less is known about the degree to which the dorsal visual pathway reorganizes following damage in childhood. Some researchers have hypothesized that the dorsal pathway may be particularly vulnerable to disruption in childhood relative to the ventral pathway – resulting in permanent and long-lasting deficits (Braddick, Atkinson, & Wattam-Bell, 2003; Grinter, Maybery, & Badcock, 2010). Indeed, the developmental disorders that most commonly cause visual impairments in children are those that affect dorsal processing (Flanagan, Jackson, & Hill, 2003). These include cerebral visual impairment (CVI), Fragile X syndrome, William’s syndrome, and cerebral palsy (Grinter et al., 2010; Macintyre-Be on et al., 2010). These children demonstrate poor performance on tasks dependent on the dorsal pathway, such as global motion perception (Jakobson, Frisk, & Downie, 2006), motor coordination (Hocking, Bradshaw, & Rinehart, 2008), and visuospatial processing (Bellugi, Lichtenberger, Jones, Lai, & St. George, 2000; Cornish, Munir, & Cross, 1998, 1999), while demonstrating spared performance on tasks linked to the ventral pathway, such as visual object recognition.

One possible explanation for the greater vulnerability of the dorsal pathway is that it matures earlier than the ventral pathway and, therefore, has a smaller window of plasticity – making it less resilient to disruption. Indeed, studies with typically developing human and non-human primates have shown that the anatomical cytoarchitecture of PPC is more adult-like than the VOTC in infancy and early childhood (Bourne & Rosa, 2006; Ciesielski et al., 2019; Distler, Bachevalier, Kennedy, Mishkin, & Ungerleider, 1996). Furthermore, human and monkey neonates exhibit more mature magnocellular than parvocellular processing (Hammarrenger et al., 2003; Kogan, Zangenehpour, & Chaudhuri, 2000; Rakic, Barlow, & Gaze, 1977), which are the primary subcortical inputs to dorsal and ventral pathways, respectively. Thus, because the structure of the dorsal pathway is in place, or stable, earlier in development than the ventral pathway, it may be less amenable to reorganization following disruption. However, comparisons between ventral and dorsal pathways are difficult to measure in most pediatric patient populations because there are many patient-specific factors that influence how well a patient recovers, such as the extent of damage, etiology of the disease, age of onset of damage as well as age at the time of surgery.

One recent study sought to overcome some of these limitations by testing a patient with damage to both dorsal and ventral pathways. Specifically, Ahmad, Behrmann, Patterson, and Freud (2022) conducted a study with patient TC who had areas of both her left PPC and VOTC surgically removed to treat drug-resistant epilepsy. They found that TC was impaired on tasks that required dorsal pathway processing, namely, grasping a block with her fingers, but not tasks that required ventral pathway processing, namely, pantomiming or visually matching the size of a block using her fingers. Thus, although TC’s left dorsal and ventral pathways were removed concurrently, these findings suggest that functions supported by the ventral pathway recovered, whereas those supported by the dorsal pathway did not. Importantly, because they tested dorsal and ventral capacities in a single patient with damage to both pathways, they were able to rule out alternative explanations related to age at time of surgery and other aspects of disease etiology.

Although TC’s pattern of deficit suggests that ventral and dorsal pathways showed differential capacity for reorganization, there remain several open questions. For instance, it is unclear whether the two pathways were equally damaged, and, thus, whether TC’s intact visual matching abilities reflects greater sparing of VOTC tissue relative to PPC. Furthermore, it is unclear to what degree each task involved lateralized processing or could be accomplished with either hemisphere. For instance, the grasping task may have required relatively more input from the left hemisphere compared to the visual matching task. Indeed, it is unclear to what degree the visual matching is lateralized to one hemisphere. In other words, visual matching might not rely on left VOTC as much as grasping relies on left PPC. Thus, one explanation for TC’s performance on the visual matching task is not that the capacities of the ventral pathway recovered, but that her visual matching abilities were only minimally disrupted in the first place. By contrast, because the grasping task is typically supported by the contralateral hemisphere, a greater degree of reorganization would be needed to recover normal grasping abilities.

In the current study, we sought to overcome these methodological challenges by testing patients with roughly comparable damage to both ventral and dorsal pathways, and by using tasks that are known to elicit lateralized processing in each pathway. Specifically, we conducted functional magnetic resonance imaging (fMRI) with patients who had undergone childhood hemispherectomy surgery – a removal or disconnection of an entire hemisphere – while they completed functional localizer tasks designed to elicit lateralized responses from left or right ventral or dorsal pathways. These included localizers for word and face regions, which are known to elicit greater responses in left and right VOTC, respectively (Behrmann & Plaut, 2020), as well as localizers for tool and object global form regions, which are known to elicit greater responses in the left and right PPC, respectively (Ayzenberg & Behrmann, 2022; Garcea & Mahon, 2014).

Hemispherectomy, rather than lobectomy or laser ablation, is undertaken in individuals whose epilepsy typically has multiple foci and leaves little to no cortical tissue in the resected hemisphere (see Figure 1). Remarkably, despite the size of the lesion, these patients generally have good post-surgical cognitive (Devlin et al., 2003; Pulsifer et al., 2004) and visual (Koenraads et al., 2014) outcomes, and, in many cases, show cognitive improvements, especially if the surgery is early in childhood (Helmstaedter et al., 2020). Importantly, these patients can learn to read (Danelli et al., 2013) and there are few clinical reports of prosopagnosia or object agnosia post-surgery. These findings suggest that the functions of the ventral pathway may have reorganized to the intact contralateral hemisphere.

**Figure 1.**
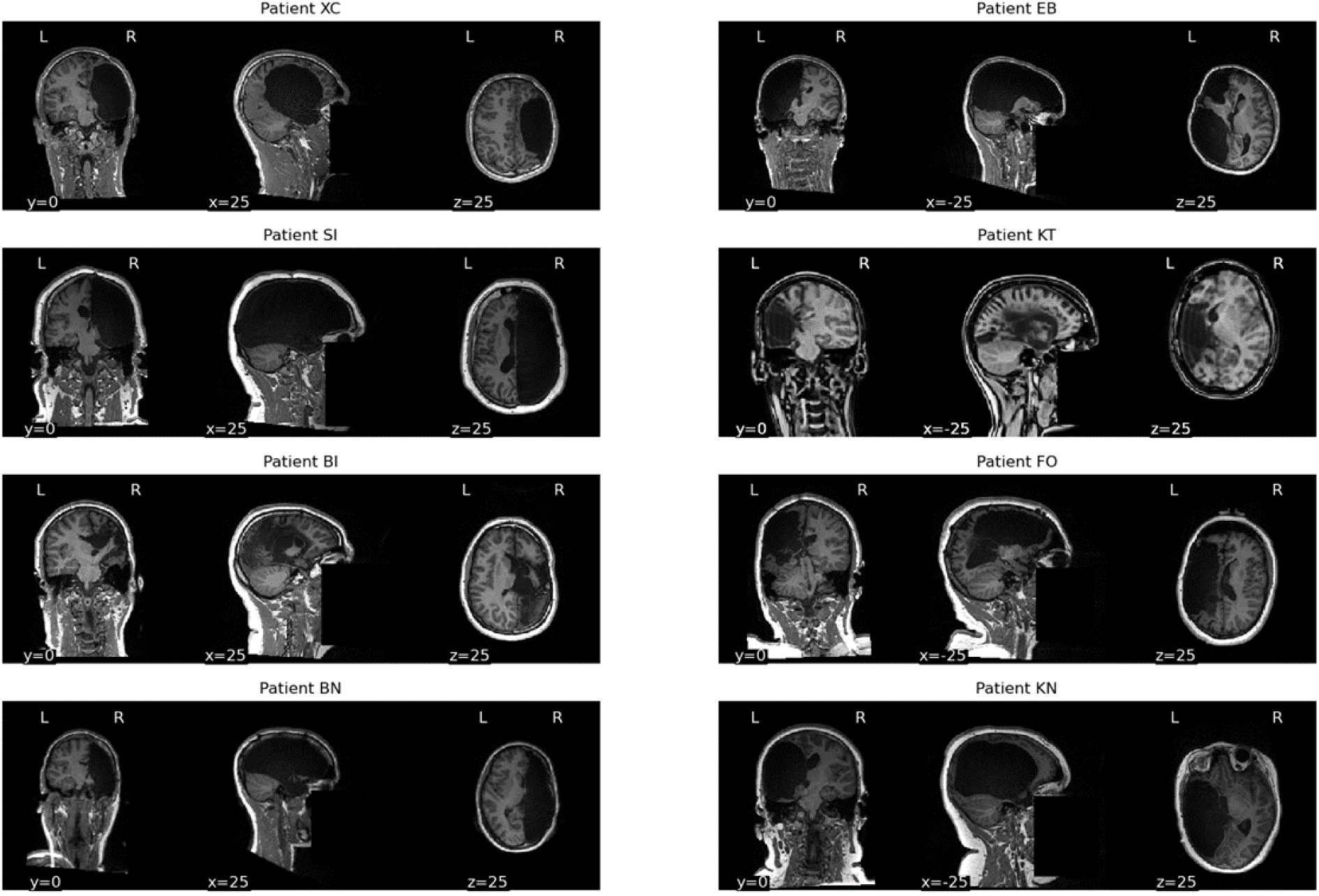
Anatomical MRI images illustrating the intact and resected portions of each patient’s brain. Images have been defaced to protect the identity of each patient.

However, few studies have directly examined the degree to which both ventral and dorsal visual pathways reorganize in the same individual, particularly following an extensive resection as in hemispherectomy. Thus, by testing a patient population with widespread damage to ventral and dorsal pathways and using tasks that are known to elicit lateralized processing in healthy participants, we can directly test the capacity of the visual system to reorganize and, in doing so, can shed light on the developmental trajectory of each pathway. To foreshadow our results, overall, we found that a greater number of hemispherectomy patients showed reorganization in the ventral pathway than the dorsal pathway, supporting the hypothesis that the dorsal pathway mature earlier than the ventral pathway.

## Materials and Methods

### 2.1 Participants

Eight patients who had undergone hemispherectomy in childhood were recruited (*M*_age_ = 19.38, Range: 12-37 years) either from the Pediatric Epilepsy Surgery Program at University of Pittsburgh Medical Center Children’s Hospital of Pittsburgh or the Pediatric Epilepsy Surgical Alliance. Of these, four patients had right hemispherectomies (patients: XC, SI, BI, BN) and four had left hemispherectomies (patients: EB, KT, FO, KN). Each patient completed four localizer tasks designed to elicit a lateralized response in the left or right ventral and dorsal pathways: words (left ventral), faces (right ventral), tools (left dorsal), and global form (right dorsal), with the exception of patient KT who was unable to complete the tool localizer. All patients had normal or corrected-to-normal vision in their intact hemifield. For specific patient ages and surgery information, see Table 1.

**Table 1.** Patient demographic and surgery information. Because several patients underwent revision surgeries, the ‘age at surgery’ column depicts their age at their most recent surgery. Ages are presented in years.

| Patient | Sex | Age at testing | Age at surgery | Surgery Type | Intact Hemisphere |
| --- | --- | --- | --- | --- | --- |
| XC | M | 17 | 1.0 | Right anatomical hemispherectomy | Left |
| SI | M | 37 | 11 | Right anatomical hemispherectomy | Left |
| BI | F | 16 | 1.3 | Right hemispherotomy | Left |
| BN | F | 18 | 9 | Right anatomical hemispherectomy | Left |
| EB | F | 16 | 1.1 | Left anatomical hemispherectomy | Right |
| KT | F | 15 | 13 | Left functional hemispherectomy | Right |
| FO | F | 19 | 4 | Left anatomical hemispherectomy | Right |
| KN | M | 12 | 1.7 | Left anatomical hemispherectomy | Right |

We also tested 44 control participants (*M*_age_ = 22.53, Range: 13-38 years). Fifteen control participants completed the word and face localizer task, and 18 completed the tool and global form localizer task. These control participants were recruited as part of other ongoing projects (Ayzenberg & Behrmann, 2022; Liu et al., 2019; Maallo et al., 2020). Six additional participants completed all four tasks. All control participants were right-handed and had normal or corrected-to-normal visual acuity.

All participants and/or their guardians gave informed consent and assent according to a protocol approved by the Institutional Review Board (IRB) of Carnegie Mellon University and the University of Pittsburgh, and received payment for their participation.

#### Brief case description

All patients recruited for this study were seizure free at the time of testing and were capable of both reading and writing. Patients did not have a clinical history of alexia, prosopagnosia, or object agnosia. Behavioral pre-testing further revealed that patients showed good performance on word, face, and shape recognition tasks, with many patients (though not all) performing as well as controls (see Supplemental Figures 1 and 2 for results for each patient). Patients are hemiplegic with impaired motor control of limbs contralateral to the resected hemisphere while limbs contralateral to their intact hemisphere appear to be intact. Indeed, patients successfully use their hand for fine motor skills like writing or manipulating cutlery to eat. Thus, under gross observation, patients’ behavioral profiles show minimal, if any, evidence of word, face, tool use, or shape perception deficits.

### 2.2 MRI scan parameters and analysis

Scanning was done on a 3T Siemens Prisma scanner at the CMU-Pitt Brain Imaging Data Generation & Education (BRIDGE) Center. Whole-brain functional images were acquired using a 64-channel head matrix coil and a gradient echo single-shot echoplanar imaging sequence. Whole-brain, high-resolution T1-weighted anatomical images (repetition time = 2300 ms; echo time = 2.03 ms; voxel size = 1 × 1 ×1 mm) were also acquired for each participant to register of the functional images into a common space. The acquisition protocol for each functional run of the word and face localizer consisted of 69 slices, repetition time = 2 s; echo time = 30 ms; flip angle = 79°; voxel size = 3 × 3 × 3 mm, multi-band acceleration factor = 3. The acquisition protocol for each functional run of the tool and global form localizer consisted of 48 slices, repetition time = 1 s; echo time = 30 ms; flip angle = 64°; voxel size = 3 × 3 × 3 mm, multi-band acceleration factor = 4.

All images were skull-stripped (Smith, 2002) and registered to participants’ native anatomical space. Prior to statistical analyses, images were motion corrected, de-trended, and intensity normalized. An additional 18 motion regressors generated by FSL were also included. All data were fit with a general linear model consisting of covariates that were convolved with a double-gamma function to approximate the hemodynamic response function. Analyses were conducted using FSL (Smith et al., 2004), and the nilearn and nibabel packages for Python (Abraham et al., 2014).

### 2.3 Localizer tasks

We administered four localizer tasks designed to elicit a lateralized response in the left or right ventral and dorsal pathways: words (left ventral), faces (right ventral), tools (left dorsal), and global form (right dorsal). Whenever possible, we collected three runs of each localizer task. However, due to participant tolerance and time constraints, some patient and control participants were only able to complete two runs of some tasks. Thus, because each participant contributed at least two runs of a given localizer, we restricted all of our analysis to just two runs of data.

#### Word and face localizer

On each run of the word and face localizer (378 s), participants viewed blocks of images of words, faces, objects, houses, or box-scrambled images (Figure 2A and 2B). Each block contained 16 images displaying 15 unique instances, with 1 repeat stimulus randomly inserted per sequence. All stimuli subtended ∼4° visual angle on screen. Each image was presented for 800 ms with a 200 ms interstimulus interval (ISI) for a total of 16 s per block. The image order within the block was randomized. Participants viewed 3 repetitions of each block per run in a pre-determined random sequence used for all participants. To maintain attention, participants performed a one-back task, responding to the repetition of an image on consecutive presentations. Word representations were measured as a greater response to the word condition than the object condition. Similarly, face representations were measured as a greater response to the face condition than the object condition.

**Figure 2.**
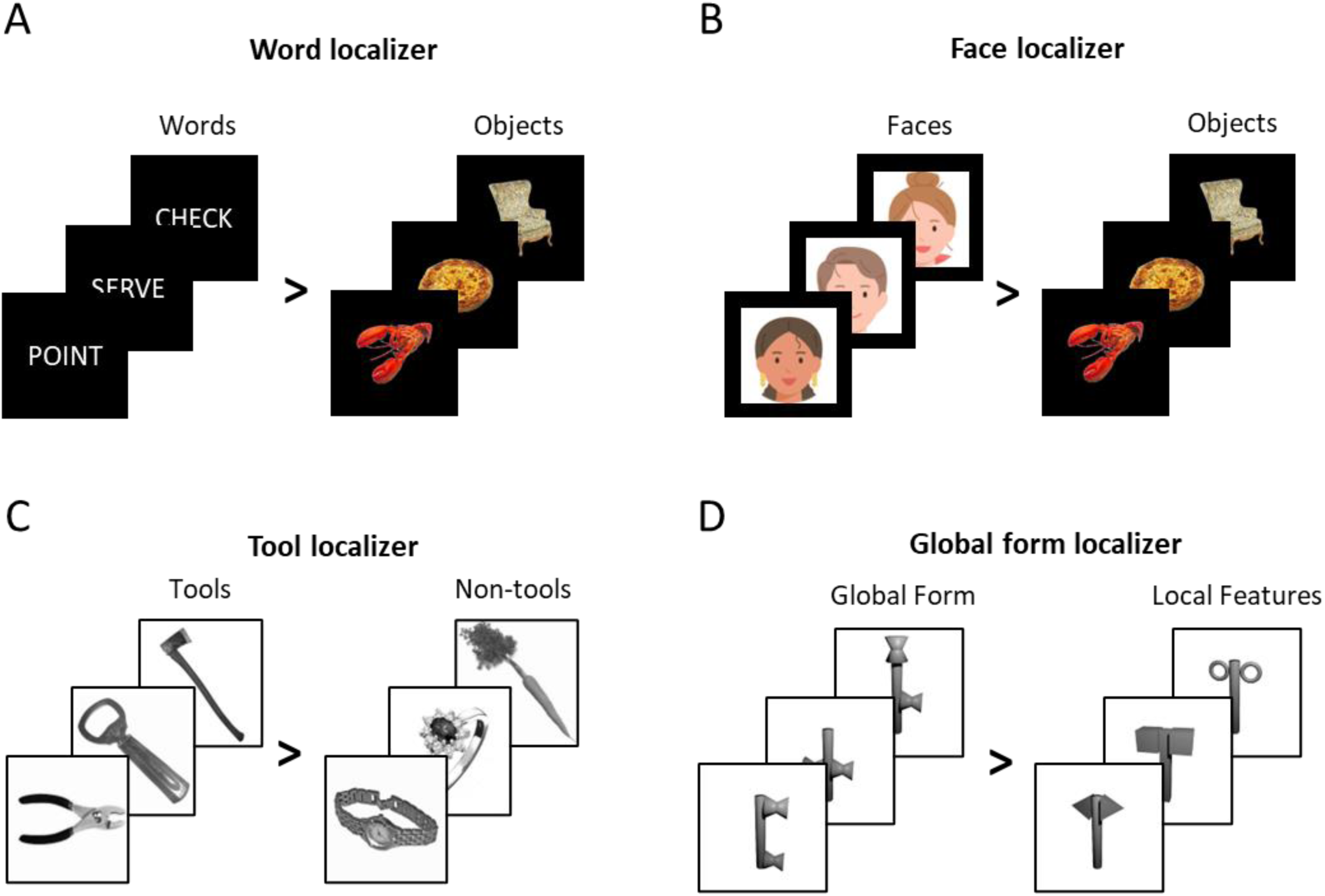
Example stimuli from the (A) word, (B) face, (C) tool, and (D) global form localizers. Note, in this example, face photographs were replaced with illustrations to comply with bioRxiv’s submission policy.

#### Tool localizer

On each run of the tool localizer (340 s), participants viewed blocks of object images that contained tools (tool condition), manipulable non-tool objects (non-tool condition), or box-scrambled object images (scrambled conditions; Figure 2C). Following previous work (Mahon et al., 2007), we defined tools here as manipulable objects whose physical form is directly related to their function (e.g., a hammer). By contrast, manipulable non-tool objects are those that can be arbitrarily manipulated, but whose form is not directly related to their function (e.g., a carrot). Each condition was comprised of ten instances each of tools, non-tools, or scrambled object images (Chen, Garcea, Jacobs, & Mahon, 2018; Chen, Garcea, & Mahon, 2016). Each block contained 20 images, displaying each possible tool, non-tool, or scrambled image twice per block. All stimuli subtended ∼6° visual angle on screen. Each image was presented for 700 ms with a 100 ms interstimulus interval (ISI) for a total of 16 s per block. The image order within the block was randomized. Participants also viewed blocks of fixation (16 s). Participants viewed 5 repetitions of each block per run, with blocks presented in a pseudorandom order under the constraint that all four block types (tool, non-tool, scrambled, fixation) were presented once before repetition. To maintain attention, participants performed an orthogonal one-back task, responding to the repetition of an image on consecutive presentations. Tool representations were measured as those voxels that responded more to the tool than the non-tool condition.

#### Global form localizer

On each run of the global form localizer (320 s), participants viewed blocks of object images in which either the spatial arrangement of component parts varied from image to image (global form condition), while the parts themselves stayed the same; or the features of the component parts varied from image to image (local feature condition), while the spatial arrangement of the parts stayed the same (Figure 2D). Objects could have one of 10 possible spatial arrangements, and one of 10 possible part features. Spatial arrangements were selected to be qualitatively different from one another as outlined by the recognition-by-components (RBC) model (e.g., end-to-end; end-to-middle; Biederman, 1987). The component parts were comprised of qualitatively different features as outlined by the RBC model (e.g., sphere, cube). Because many dorsal regions are particularly sensitive to an object’s orientation and axis of elongation (Sakata et al., 1998), all objects were presented in the same orientations and were organized around the same elongated segment, ensuring that they have identical principal axes. Stimuli subtended ∼6° visual angle on screen.

Each block of the global form localizer contained 20 images, displaying each spatial arrangement or part feature twice per block depending on the condition. Each image was presented for 800 ms with a 200 ms interstimulus interval (ISI) for a total of 20 s per block. To minimize visual adaptation, the location of object images on the screen varied by ∼2° every trial. The image order within the block was randomized. Participants also viewed blocks of a fixation cross (20 s). Participants viewed 5 repetitions of each block per run, with blocks presented in a pseudorandom order under the constraint that all three block types (relations, feature, fixation) were presented once before repetition. To maintain attention, participants performed an orthogonal one-back task, in which they responded via key press when detecting the repetition of an image on consecutive presentations. Global form representations were quantified as those voxels that responded more to the global form than the local feature condition.

### 2.4 Statistical Analysis

#### Neural response

We first measured whether patients demonstrated any statistically reliable neural responses to the conditions of interest. We created large-scale ventral and dorsal region-of-interest (ROI) binary masks using probabilistic parcels (Julian, Fedorenko, Webster, & Kanwisher, 2012; Wang, Mruczek, Arcaro, & Kastner, 2014). For the ventral visual pathway, we included parcels beginning at visual area 4 (V4) and ending at anterior portions of the fusiform, encompassing the typical positions of the word area, namely the VWFA, as well as face areas, namely the occipital face area (OFA) and FFA (see Supplemental Figure 3). For the dorsal visual pathway, we included parcels beginning at visual area 3 A/B (V3A/B) and ending at intraparietal sulcus area 5 (IPS5) and the superior parietal lobe (SPL) (see Supplemental Figure 3). Ventral and dorsal ROIs were purposefully created to encompass a large portion of cortex so as to accommodate patients’ potentially altered anatomy and, thus, to ensure that we captured any statistically reliable neural responses in their ventral and dorsal pathways. However, an ROI-free analysis examining the entire hemisphere was also performed.

Because the patients’ overall anatomy is altered due to their missing hemisphere, conventional registration techniques are not always successful. Thus, to register each ROI from MNI standard space to each individual patient, we first created a mirror symmetric version of each patient’s brain by combining the anatomical image of the intact hemisphere with a mirror-flipped version of the intact hemisphere. We then computed the registration transformation between the MNI anatomical template and each patient’s mirror symmetric anatomical image. This final transformation matrix was then used to register the ventral and dorsal ROIs to each patient’s preserved hemisphere (see Supplemental Figures 3). Control data were registered using standard procedures.

Neural responses within ventral and dorsal masks were measured as those voxels that survived a liberal uncorrected threshold of *p* < .01. We used a lax threshold because only a limited amount of data was collected for each participant (two runs per localizer), thereby limiting the statistical power. Moreover, because we were ultimately interested in comparisons between patient and control groups, it was primarily important that the same threshold was used for each participant. Results were qualitatively the same at higher thresholds, as well as when a threshold-free analysis was performed using all positive voxel values.

#### Anatomical location

To test whether the anatomical locations of responses within ventral and dorsal pathways aligned with those of controls, for each condition, we evaluated the distance between the peak response in the patients and the peak response in the controls.

To do so, we registered all participants (patients and controls) to MNI space and then computed the coordinate of the peak response to each condition within ventral and dorsal masks (ventral: words and face; dorsal: tools and global form). Because prior work has shown that the neural response to faces and global form (Ayzenberg & Behrmann, 2022; Kamps, Morris, & Dilks, 2019) typically show both a posterior (faces: OFA; global form: posterior IPS) and anterior (faces: FFA; global form: anterior IPS) cluster, we further split the analysis for these conditions into posterior and anterior regions.

As there were relatively few patients compared to controls, we conducted our analyses on a patient-by-patient basis using non-parametric statistics. Specifically, for every condition (word, face, tool, global form), hemisphere (left, right), and region (ventral, dorsal), we computed bootstrapped 95% confidence intervals using the control data. On every resample of the data, 4 control participants (to match the number of patients with each hemisphere) were randomly selected (without replacement), and the mean Euclidean distance between their peak response for a condition and the remaining controls was calculated. This procedure was then repeated 10,000 times, thereby creating a distribution of distance values. We then tested whether the distance for each individual patient fell below the control distribution.

#### Selectivity

We measured whether patients exhibited normal levels of selectivity for each stimulus condition by computing the mean activation to each localizer contrast, the mean active volume, and a composite score known as summed selectivity (Vin, Blauch, Plaut, & Behrmann, 2023). Although mean activation and total active volume are common measures of selectivity, they only provide partial insight into the neural response profile for a given condition. For instance, the mean activation amplitude for the words > objects contrast may be normal for a patient relative to controls, but the patient may exhibit a much smaller area of activation compared to controls. By contrast, the overall area of activation in a patient may be comparable to that of controls, but the mean activation may be overall lower. Summed selectivity sums each significant voxel value, thereby providing a holistic measure that captures both overall activation strength and the total active area.

For word and face conditions, we computed selectivity metrics within broad ventral binary masks (Julian et al., 2012; Wang et al., 2014) and selecting those voxels that survived a liberal uncorrected threshold of *p* < .01. Similarly, for tools and global form, we computed selectivity metrics within broad dorsal binary masks (Wang et al., 2014) and selecting those voxels that survived a liberal uncorrected threshold of *p* < .01.

Mean activation was computed as the mean of all standardized parameter estimate values (betas) above this threshold. Active volume (mm^3^) was computed as a count of the total number of voxels above threshold then cubed. Finally, summed selectivity was computed by summing the standardized parameter estimate values for each surviving voxel. Because summed selectivity is influenced by the total number of voxels within a region and because participants have different sized brains, we normalized summed selectivity values for each condition by the total number of available voxels within each participants’ ROI mask for that condition. For these values to be more easily interpretable, we rescaled them by a factor of 1000. As above, we conducted our analyses on a patient-by-patient basis using non-parametric statistics. Each patient’s summed selectivity value was compared to bootstrapped 95% confidence using the control data (10,000 resamples with replacement).

#### Decoding

Even if patients exhibited abnormal selectivity for each stimulus condition, the distributed pattern of their neural response may still potentially support encoding of each stimulus category. Thus, we also tested whether we could decode each stimulus condition from the patients’ multivariate neural responses. We extracted the multivariate neural response (averaged across time) for each block of trials for a particular stimulus condition (word, face, tool, and global form extracted from blocks of each localizer) from each region (left, right hemisphere; ventral, dorsal pathway). Then, using a 30-fold cross-validation procedure, a Support Vector Machine (SVM) classifier was trained on the multivariate pattern for 80% of the blocks, and then tested on the held-out 20%. Decoding for each condition was tested against the multivariate neural response of its contrast (words vs. objects; faces vs. objects; tools vs. non-tools; global form vs. local features).

## Results

### 3.1 Preliminary Analyses

We first examined what proportion of control participants had significant activation to each condition (word, face, tool, global form) within each condition’s preferred hemisphere (left, right) and ROI (ventral, dorsal). We found that the majority of control participants who completed the ventral localizer tasks exhibited significant activation to words in their left VOTC (20/21 participants) and faces in their right VOTC (20/21 participants). Similarly, every control participant who completed the dorsal localizer tasks exhibited significant activation to tools in their left dorsal pathway (24/24) and global form in their right dorsal pathway (24/24).

Next, we examined whether the number and location of significant clusters for each condition in the controls corresponded to their typical location based on the literature. Examination of the group activation maps in the ventral pathway revealed one posterior ROI in the ventral pathway for words, corresponding to the VWFA (see Figure 3B; Dehaene & Cohen, 2011) and two ROIs for faces corresponding to OFA and FFA (see Figure 4B; Kanwisher, McDermott, & Chun, 1997; Pitcher, Walsh, & Duchaine, 2011). In the dorsal pathway, we observed one anterior ROI for tools, corresponding to the superior parietal lobule (see Figure 5B; Johnson-Frey, 2004), and two ROIs for global form, corresponding to posterior and anterior IPS (see Figure 6B; Ayzenberg & Behrmann, 2022).

**Figure 3.**
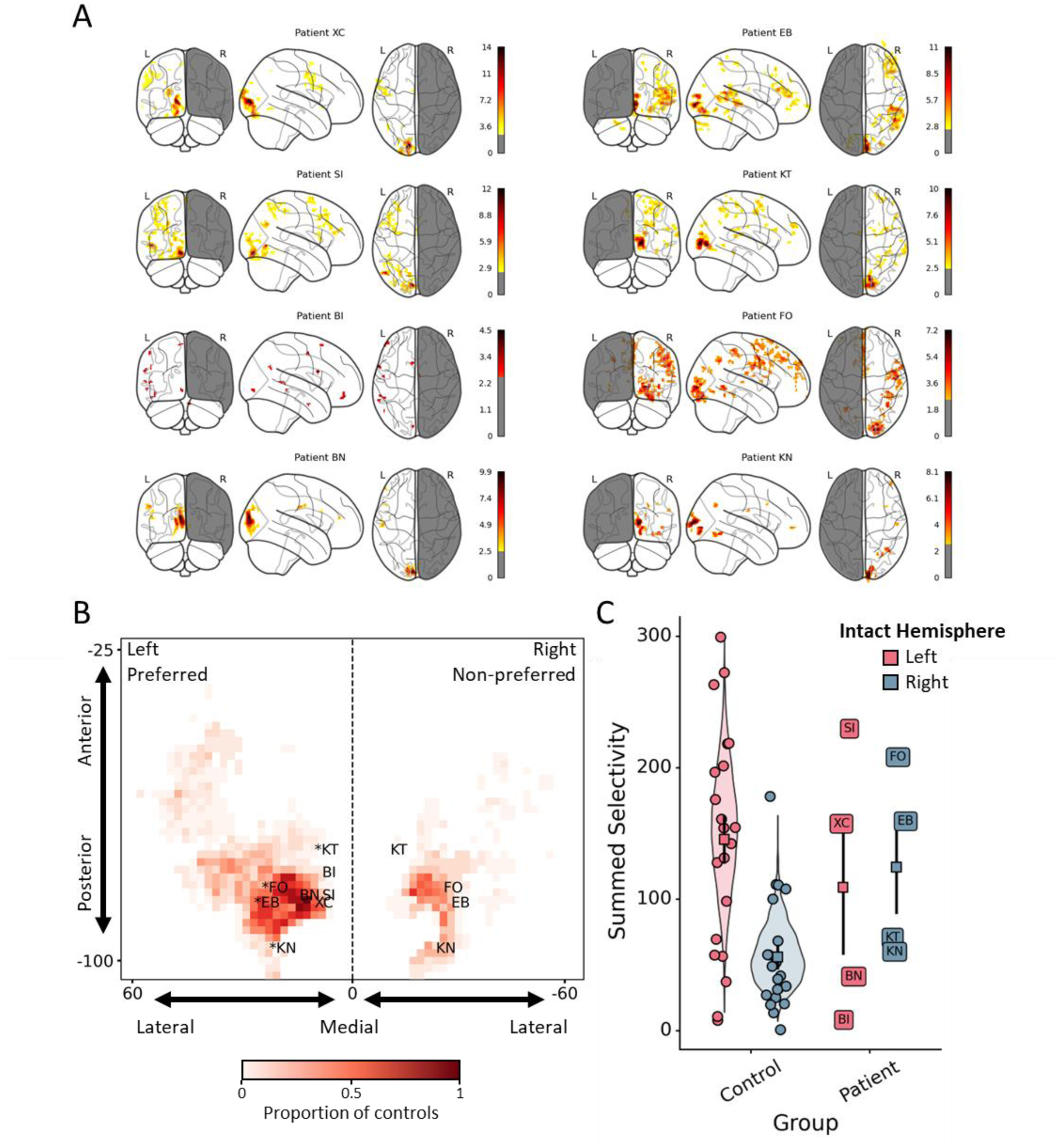
Results from the word localizer. (A) Whole brain responses to words > objects in patients with intact left or right hemispheres. (B) Visualization of word responses in controls and patients in the ventral pathway. The heatmap illustrates the 2D spatial distribution of group responses to words in controls, with darker colors indicating that a larger proportion of controls had significant activation at that coordinate. Each labeled point refers to the peak coordinate for a patient. Patients with an intact right hemisphere have been projected onto the preferred hemisphere of the map and are marked with an asterisk (*). (C) Summed selectivity for words in the ventral pathway. Violin plots depict the bootstrapped distribution of control participants’ scores. Each unlabeled point refers to a single control participant’s data, and each labeled point corresponds to a single patient. Square points and error bars refer to the mean and standard error of each group, respectively.

**Figure 4.**
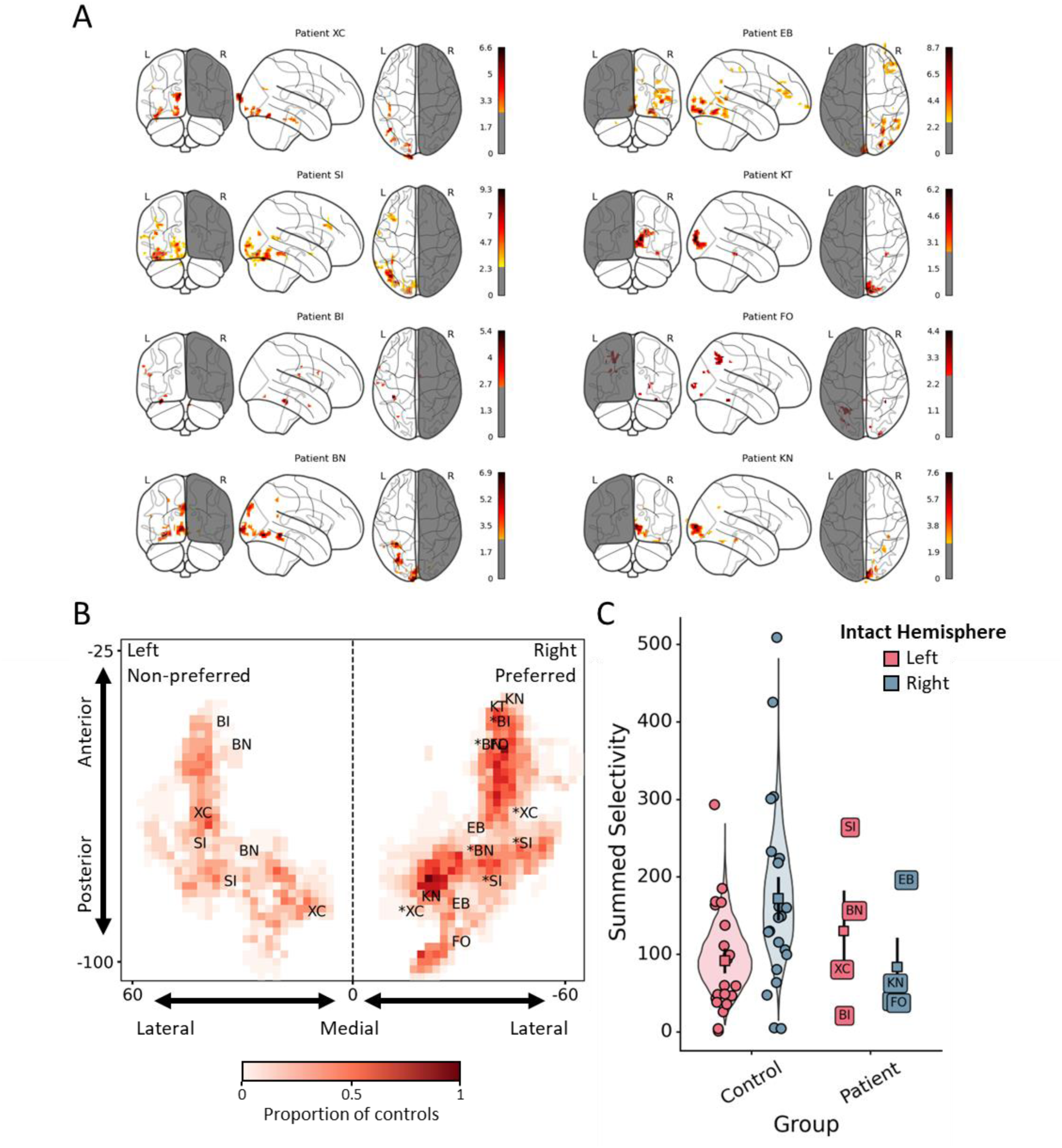
Results from the face localizer. (A) Whole brain responses to faces > objects in patients with intact left or right hemispheres. (B) Visualization of face responses in controls and patients in the ventral pathway. The heatmap illustrates the 2D spatial distribution of group responses to words in controls, with darker colors indicating that a larger proportion of controls had significant activation at that coordinate. Each labeled point refers to the peak coordinate for a patient. Patients with an intact left hemisphere have been projected onto the preferred hemisphere of the map and are marked with a *. (C) Summed selectivity for faces in the ventral pathway. Violin plots depict the bootstrapped distribution of control participants’ scores. Each unlabeled point refers to a single control participant’s data, and each labeled point corresponds to a single patient. Square points and error bar refer to the mean and standard error of each group, respectively.

**Figure 5.**
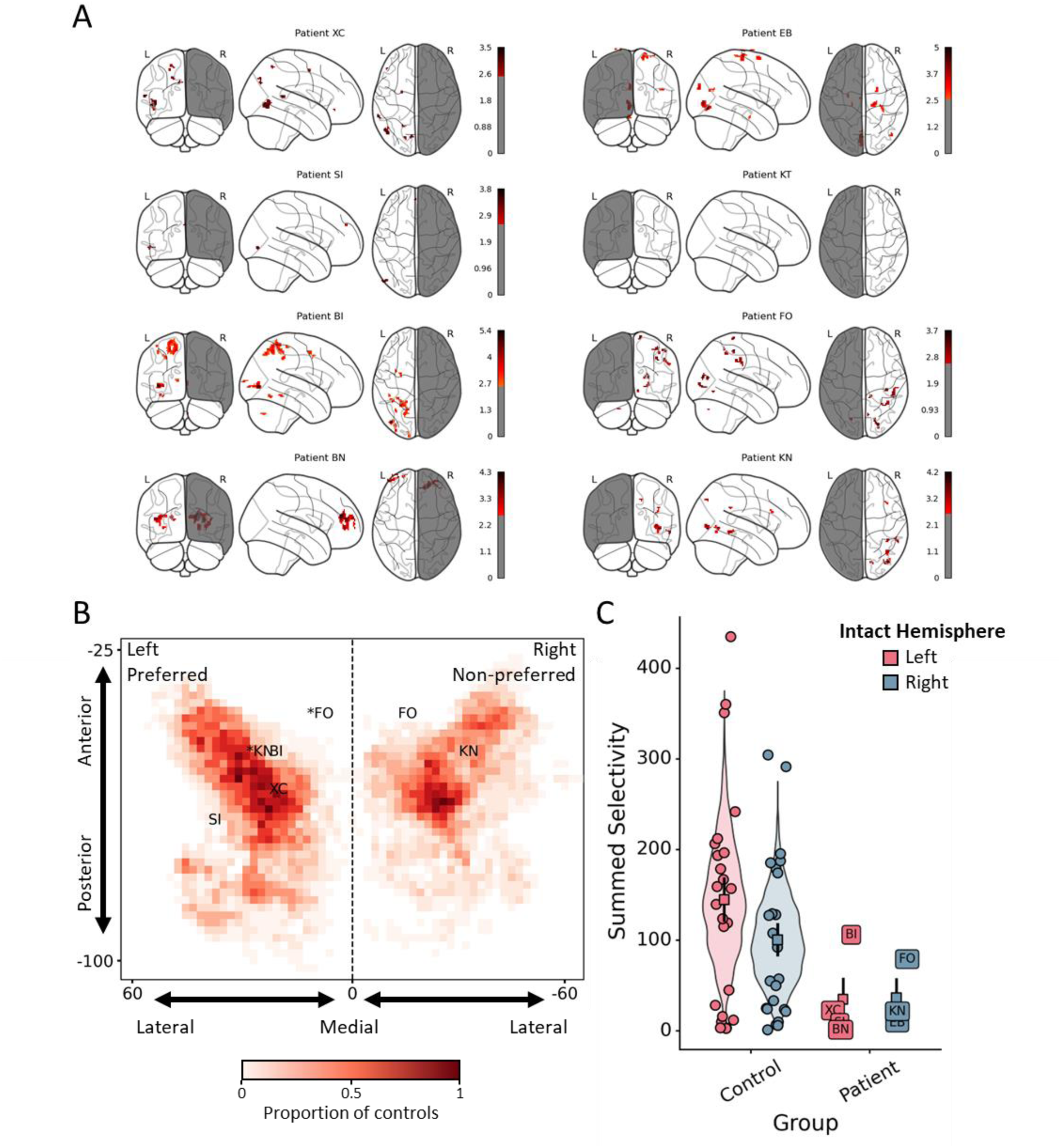
Results from the tool localizer. (A) Whole brain responses to tools > non-tools in patients with intact left or right hemispheres. (B) Visualization of tool responses in controls and patients in the dorsal pathway. The heatmap illustrates the 2D spatial distribution of group responses to tools in controls, with darker colors indicating that a larger proportion of controls had significant activation at that coordinate. Each labeled point refers to the peak coordinate for a patient. Patients with an intact right hemisphere have been projected onto the preferred hemisphere of the map and are marked with a *. (C) Summed selectivity for tools in the dorsal pathway. Violin plots depict the bootstrapped distribution of control participants’ scores. Each unlabeled point refers to a single control participant’s data, and each labeled point corresponds to a single patient. Square points and error bar refer to the mean and standard error of each group, respectively.

**Figure 6.**
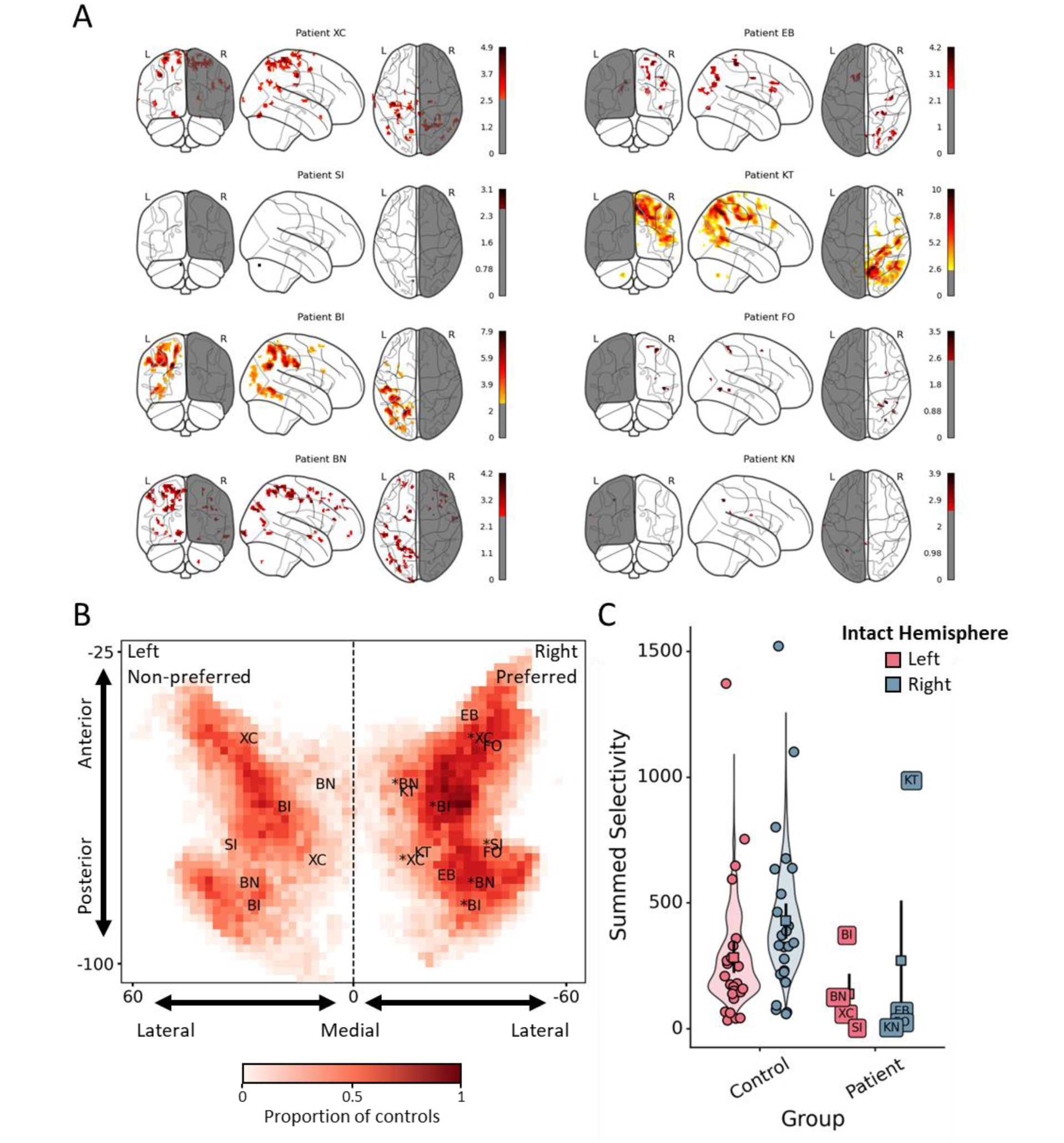
Results from the global form localizer. (A) Whole brain responses to global form > local features in patients with intact left or right hemispheres. (B) Visualization of global form responses in controls and patients in the dorsal pathway. The heatmap illustrates the 2D spatial distribution of group responses to words in controls, with darker colors indicating that a larger proportion of controls had significant activation at that coordinate. Each labeled point refers to the peak coordinate for a patient. Patients with an intact left hemisphere have been projected onto the preferred hemisphere of the map and are marked with a *. (C) Summed selectivity for global form in the dorsal pathway. Violin plots depict the bootstrapped distribution of control participants’ scores. Each unlabeled point refers to a single control participant’s data, and each labeled point corresponds to a single patient. Square points and error bar refer to the mean and standard error of each group, respectively.

Finally, we tested whether control participants showed the predicted pattern of lateralization for each condition. In the ventral pathway, this analysis revealed stronger summed selectivity for words in the left hemisphere compared to the right, *t*(20) = 6.18, *p* < .001, *d* = 1.34 (Figure 3C) and stronger summed selectivity for faces in the right hemisphere compared to the left, *t*(20) = 4.45, *p* < .001, *d* = 0.75 (see Figure 4C). Similarly, in the dorsal pathway, we found stronger summed selectivity for tools in the left hemisphere compared to the right, *t*(23) = 3.19, *p* = .004, *d* = 0.41 (see Figure 5C), and stronger summed selectivity to global form in the right hemisphere compared to the left, *t*(23) = 5.83, *p* < .001, *d* = 0.45 (see Figure 6C). Altogether these analyses replicate the previously reported response profiles for each condition.

### 3.2 Patient Analyses

Our primary question of interest is whether functions that are typically activated to a greater extent in one hemisphere, as demonstrated in the control data, reorganize to the contralateral hemisphere following hemispherectomy. To this end, we focused on analyses comparing the response of each patient’s intact hemisphere to the ‘preferred’ (typical localization) hemisphere for each condition in controls (words: patient right vs. control left; faces: patient left vs. control right; tools: patient right vs. control left; global form: patient left vs. control right). For the results of every possible comparison, see Table 2 and supplemental materials. As described in the methods, all patient metrics were compared to 95% CIs computed from a bootstrapped distribution of control participants. For the selectivity analyses, we focused on the summed selectivity metric as this provides an overall description of the response profile to a condition (see Methods). Moreover, we specifically tested whether patient responses were *below* the distribution of selectivity values for controls, because selectivity values within or above control distribution would be evidence of reorganization. The results are qualitatively similar when examining mean activation and total active volume (see Supplemental Figures 4-5).

**Table 2.**
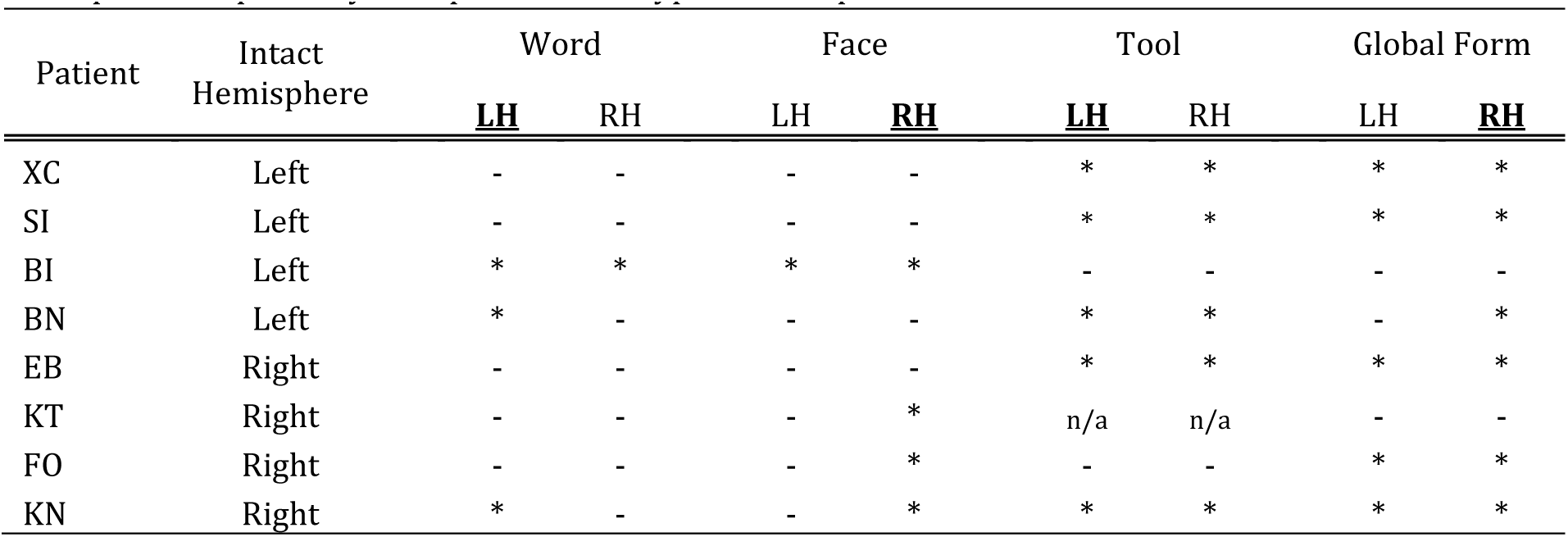
At-a-glance summary of selectivity results. Each row indicates whether a patient’s summed selectivity score was below the control distribution (95% CIs). Asterisks (*) indicate that the patient’s score was below the control distribution, dashes (-) indicate that the patient’s score was inside or above the control distribution. ‘LH and ‘RH’ indicate whether the scores were compared to controls’ left hemisphere or right hemisphere, respectively. The ‘preferred’ or typical hemisphere for each condition is underlined and in bold.

#### 3.2.1 Word representations

In controls, words are represented more strongly in the left hemisphere of VOTC than the right hemisphere. Thus, we tested whether patients with a left hemispherectomy, and, therefore, only an intact right hemisphere, demonstrated normal word representations in their right hemisphere relative to controls’ preferred left hemisphere (see Figure 3).

We first examined whether patients showed any statistically reliable responses to words in their preserved right hemisphere. This analysis revealed that all four left hemispherectomy patients showed word responses in the posterior portion of their intact right VOTC (Figure 3A-B). Next, we examined whether the location of peak responses to words in patients’ intact hemisphere aligned with controls’ peak responses in their preferred left hemisphere. This analysis revealed that none of the left hemispherectomy patients showed peak responses within the control distribution of distances (Figure 3B).

Finally, we examined whether the overall selectivity of word responses in patients was comparable to that of the controls’ preferred left hemisphere. This analysis revealed that EB, KT, and FO’s summed selectivity for words fell within the control distribution, whereas KN’s did not (see Table 2 and Figure 3C). Instead, KN’s summed selectivity to words was more comparable to control’s non-preferred right hemisphere.

Next, given that word representations are typically lateralized to the left hemisphere, one might have also predicted that patients who have an intact left hemisphere would show normal word selectivity. Our results showed that that only two of the four patients (patient SI and XC) with an intact left hemisphere showed summed selectivity that was comparable to that of controls. One other patient (patient BN) showed summed selectivity that was comparable to control’s right hemisphere. Summed selectivity for the final patient (patient BI) was lower than both the left and right hemisphere of controls. Together, these analyses provide evidence that word representations reorganize to the contralateral hemisphere following left hemispherectomy, but also, offer some evidence that hemispherectomy disrupts the typical neural organization for the intact hemisphere (see General Discussion).

#### 3.2.2 Face representations

In controls, faces are represented more strongly in the right than left hemisphere of VOTC. Thus, we tested whether patients who have had a right hemispherectomy, and, therefore, have only an intact left hemisphere, demonstrated normal face representations in their left hemisphere as compared to controls’ right hemisphere (see Figure 4).

We first examined whether patients showed any statistically reliable responses to faces in their intact left VOTC. This analysis revealed that three out of four patients, (XC, SI, and BN), exhibited face responses in the posterior portion of their intact left VOTC, and all four patients, SI, BI, and BN, exhibited face responses in the anterior portion of their intact left VOTC (see Figure 4A-B). Of these, SI and BN’s peak responses aligned with the location of controls’ peak voxel in posterior VOTC, and XC and BN’s peak response to faces aligned with the location of controls in anterior VOTC (see Figure 4B).

Finally, we examined whether the overall selectivity of faces responses in patients was comparable to controls’ preferred right hemisphere. This analysis revealed that three out of four patients’ (XC, SI, BN) summed selectivity for faces fell within the control distribution (see Table 2 and Figure 4C). Although BI showed face responses in her intact left hemisphere, they were lower than both left and right hemispheres of controls.

Next, given that face representations are typically lateralized to the right hemisphere, one might have also predicted that patients who have an intact right hemisphere would show normal face selectivity. Here, we found that only EB showed face responses that fell within the control distribution for face selectivity in the right hemisphere. However, the summed selectivity of the remaining three left hemispherectomy patients fell within the range of face responses in controls’ non-preferred left hemisphere. These analyses show evidence that face representations reorganize to the contralateral hemisphere, and also suggest that hemispherectomy may disrupt the face representations in patient’s intact right hemisphere.

#### 3.2.3 Tool representations

In controls, tools are represented more strongly in the left hemisphere of PPC than the right hemisphere. Thus, we tested whether patients who have had a left hemispherectomy, and therefore only have an intact right hemisphere, demonstrated normal tool representations in their right hemisphere as compared to control’s left hemisphere (see Figure 5).

We first examined whether patients showed any statistically reliable responses to tools in their intact right hemisphere. This analysis revealed that only FO and KN showed reliable tool responses in the anterior portion of their intact right dorsal pathway (Figure 5A-B). Next, we examined whether the location of peak responses to tools in patients’ intact hemisphere aligned with controls preferred left hemisphere. This analysis revealed only KN showed peak responses within the control distribution (Figure 5B).

Finally, we examined whether the overall selectivity of tool responses in patients was comparable to controls’ preferred left hemisphere. This analysis revealed that only FO’s summed selectivity for tools fell within the control distribution (see Table 2 and Figure 5C). Next, given that tool representations are typically lateralized to the left hemisphere, we tested whether patients who have an intact left hemisphere would show normal tool selectivity. However, our results showed that that only BI’s summed selectivity for tools fell within the control distribution. These analyses provide very little evidence that tool representations reorganize to the contralateral hemisphere, and further suggest that hemispherectomy may disrupt tool representations in patients with an intact left hemisphere.

#### 3.2.4 Global form representations

In controls, global form is represented more strongly in the right hemisphere of PPC than the left hemisphere. Thus, we tested whether patients who have had a right hemispherectomy, and therefore only have an intact left hemisphere, demonstrate normal global form representations in their left hemisphere as compared to control’s preferred right hemisphere (see Figure 6).

We first examined whether patients showed any statistically reliable responses to global form in their intact left PPC. This analysis revealed that three patients, XC, BI, and BN, exhibited global form responses in the posterior portion of their intact left PPC, and all four patients exhibited global form responses in the anterior portion of their intact left PPC (see Figure 6A-b). Of these, BI and BN’s peak responses aligned with the location of controls’ peak responses in the posterior portions of their dorsal pathway, and XC and BI’s peak responses aligned with controls in anterior portions of their dorsal pathway (see Figure 6B).

Finally, we examined whether the overall selectivity for global form in patients was comparable to controls. This analysis revealed that only BI’s summed selectivity for global form fell within the control distribution (see Table 2 and Figure 6C). Next, given that global representations are typically lateralized to the right hemisphere, we tested whether patients who have an intact right hemisphere would show normal global form selectivity. Here we found that only KT showed global form responses that were comparable to controls. These analyses show mixed evidence that global representation reorganize to the contralateral hemisphere, and some evidence that hemispherectomy impairs the global form representation of the intact right hemisphere.

#### 3.2.5 Signal Quality

One possible explanation for why patients did not show reorganization across both ventral and dorsal conditions is that they had excessive motion or poor temporal signal-to-noise ratio (tSNR). An analysis of patients’ motion in the scanner revealed that all patients (and controls) exhibited less than 0.25° of rotation and less than 0.20 mm of translation within the scanner, well within the acceptable bounds for fMRI analyses (see Supplemental Figure 6A-B). However, separate analyses of tSNR in ventral and dorsal pathways revealed that one patient’s tSNR was lower than controls in the ventral pathway (patient FO) and two patients’ tSNR lower than controls in the dorsal pathway (patients KN and FO; see Supplemental Figure 6C-D).

Although two patients showed lower than average tSNR, these findings cannot explain the presence of reorganization in the ventral pathway, but not the dorsal pathway, for the remaining patients. Furthermore, individual patients’ tSNR is largely inconsistent with their selectivity metrics. For instance, although FO was the only patient to show low tSNR for both ventral and dorsal pathways, she nevertheless showed evidence of reorganization for words in the ventral pathway, and was one of only two patients to show normal responses to tools in the dorsal pathway. By contrast, BI had the highest tSNR for the ventral pathway of all patients, and but showed little evidence of reorganization for words and faces. Similarly, XC had the highest tSNR for the dorsal pathway, but showed little evidence of reorganization in the dorsal pathway. Thus, it is unlikely that signal quality can explain our results.

#### 3.2.6 Summary

Overall, we found greater evidence of functional reorganization in the ventral pathway compared to the dorsal pathway. In the ventral pathway, six out of eight patients showed comparable summed selectivity in their intact hemisphere on par with control’s preferred contralateral hemisphere (3/4 for words; 3/4 for faces), with every patient showing at least some activation in their intact hemisphere. By contrast, a smaller proportion of patients showed reorganization in the dorsal pathway, with only two patients showing comparable summed selectivity to controls preferred contralateral hemisphere (1/3 tools; 1/4 global form).

However, we also found a number of inconsistencies, such that the preferred representation of patients’ intact hemisphere was often below the control distribution for a condition’s preferred ipsilateral hemisphere. Even across these inconsistencies, we found that the representations of patient’s ventral pathway were generally comparable to at least one hemisphere of controls, with 7 out of 8 patients showing comparable summed selectivity to controls for both words and faces. By contrast, only 2 out of 7 and 3 out of 8 showed any comparable summed selectivity for tools and global form, respectively (see Table 2). Altogether, these findings provide evidence that the ventral pathway is better able to reorganize following functional hemispherectomy than the dorsal pathway, but also that hemispherectomy may impact the preferred representations of the intact hemisphere.

### 3.3 Distributed Representations

Although patients showed little evidence of reorganization in the dorsal pathway, patients also generally show little impairment on recognition processes linked to the dorsal pathway such as tool use or shape perception. Moreover, not all patients showed normal selectivity to words and faces, and yet demonstrate strong performance on word and face perception tasks. In the absence of normal selectivity, how might patients accomplish these feats? One possibility is that the representations for each of these properties is distributed.

In the sections that follow, we provide two tests of this hypothesis by, first, examining whether summed selectivity in patients’ entire hemisphere, rather than in circumscribed regions, is comparable to controls, and second, by testing whether each condition can be decoded using the multivariate pattern of neural responses in ventral and dorsal pathways.

#### 3.3.1 Hemisphere analyses

We analyzed whether participants’ summed selectivity for each condition across their entire hemisphere is comparable to controls’ summed selectivity in a hemisphere (see Figure 7). Because we are most interested in examining reorganization, we focus our analyses on comparisons between each patients’ intact hemisphere and the preferred hemisphere for a condition in controls (words: patient right vs. control left; faces: patient left vs. control right; tools: patient right vs. control left; global form: patient left vs. control right). For the results of all patients see Supplemental Table 1.

**Figure 7.**
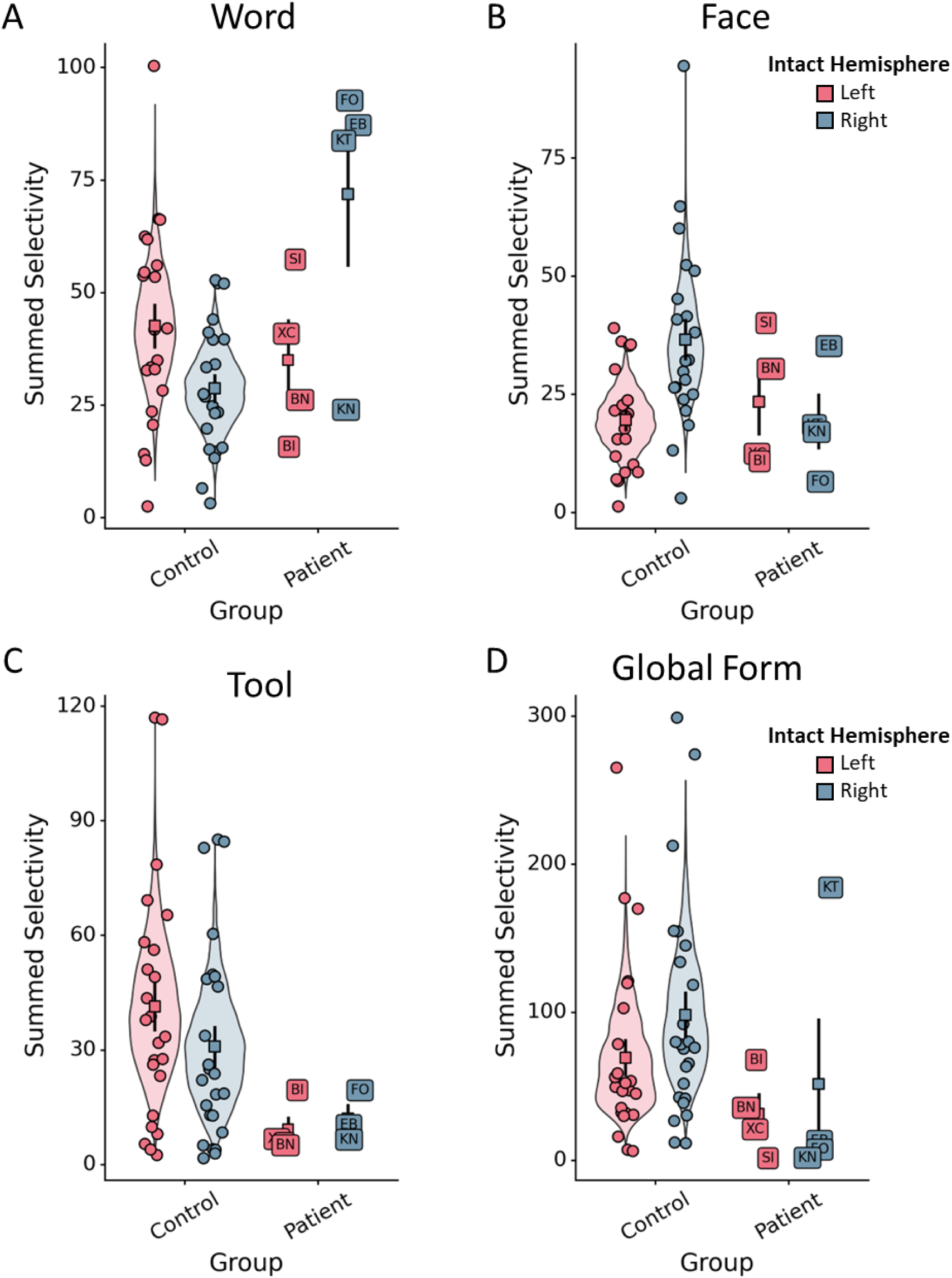
Summed selectivity for (A) words, (B) faces, (C) tools, and (D) global form in each patient and control’s hemisphere. Violin plots depict the bootstrapped distribution of control participants’ scores. Each unlabeled point refers to a single control participant’s data, and each labeled point corresponds to a single patient. Square points and error bar refer to the mean and standard error of each group, respectively.

**Figure 8.**
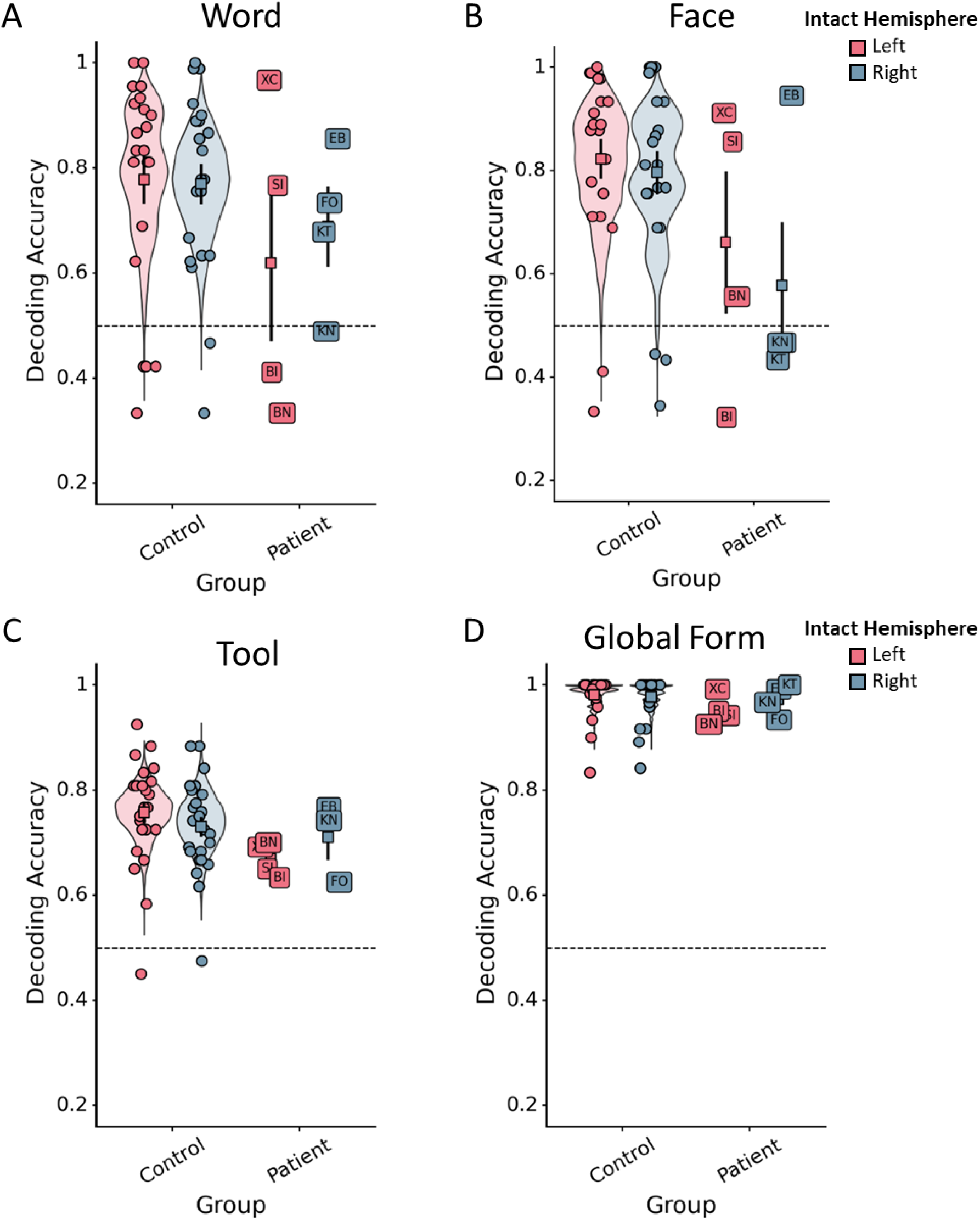
Decoding accuracy for (A) words vs. objects, (B) faces vs. objects, (C) tools vs. non-tools, and (D) global form vs. local features in each patient and control’s intact hemisphere. Violin plots depict the bootstrapped distribution of control participants’ scores. Each unlabeled point refers to a single control participant’s data, and each labeled point corresponds to a single patient. Square points and error bar refer to the mean and standard error of each group, respectively.

For the ventral conditions, we found that all four intact right hemisphere patients exhibited normal or high summed selectivity for words (see Figure 7A), and only two patients with intact left hemispheres (patients SI and BN) exhibited normal summed selectivity for faces (see Figure 7B). Interestingly, however, the word responses of EB, KT, and FO, who had intact right hemispheres, surpassed those of the controls. For the dorsal conditions, we found that only FO exhibited normal summed selectivity for tools in their intact right hemisphere (see Figure 7C), and only BI exhibited normal summed selectivity for global form in their intact left hemisphere (see Figure 7D). Thus, the overall pattern of summed selectivity across the entire hemisphere was comparable to that revealed by the analyses restricted to just ventral and dorsal pathways.

#### 3.3.2 Multivariate decoding

Next, we analyzed how well we could decode each condition of interest relative to its localizer contrast (words vs. objects; faces vs. objects; tools vs. non-tools; global form vs. local features). Here again, we are most interested in examining reorganization, and so we focused our analyses on comparisons between each patients’ intact hemisphere and the preferred hemisphere for a condition in controls (words: patient right vs. control left; faces: patient left vs. control right; tools: patient right vs. left; global form: patient left vs. control right). For the results of all patients see Supplemental Table 2.

For the ventral conditions, we found that three of four intact right hemisphere patients exhibited normal decoding accuracy for words, and three of four intact left hemisphere patients exhibited normal decoding for faces. For the dorsal conditions, we found that two intact right hemisphere patients (out of three), EB and KN, exhibited normal decoding for tools, and three out of four intact left hemisphere patients exhibited normal decoding for global form. Altogether, these findings suggest that the multivariate response for each condition could theoretically support patients’ behavioral performance.

#### 3.3.2 Summary

Overall, our analysis of the distributed pattern of responses across the entire hemisphere mirrored the region-of-interest analyses described previously. As in the first section, a greater number of patients showed normal summed selectivity in their intact ventral pathway (3/4 for words; 2/4 for faces), than in their dorsal pathway (1/3 for tools; 1/4 for global form). Our decoding analyses largely mirrored these findings for the ventral pathway, with similar proportions decoding accuracy comparable to controls (3/4 for words) and (2/4 for faces). However, in the dorsal pathway, we found that a larger proportion of patients showed comparable decoding accuracy, than selectivity (2/3 for tools; 3/4 for global form). Thus, although the dorsal pathway showed little evidence of functional reorganization at the ROI level, the distributed pattern of dorsal responses may be sufficient to support tool and global form perception.

## General Discussion

In the current study, we sought to understand the capacity of ventral and dorsal visual pathways to functionally reorganize following a large-scale surgical resection, namely, hemispherectomy. The hypothesis was that, if the dorsal pathway matures earlier than the ventral pathway, then it may be less plastic and malleable, and therefore less able to reorganize than the ventral pathway following hemispherectomy. To test this hypothesis, we conducted fMRI scans of an equal number of left and right hemispherectomy patients while they completed localizer tasks designed to elicit lateralized responses in the left or right hemisphere of ventral or dorsal pathways. Overall, we found that a greater number of patients showed reorganization in the ventral pathway than the dorsal pathway. Importantly, because we examined ventral and dorsal reorganization in the same individual patients, these results cannot be explained by between-subjects factors such as disease etiology, age of surgery, and age at the time of testing. Together, these findings suggest that the dorsal pathway may develop earlier and exhibit a smaller window of plasticity.

Overall, six out of eight patients showed reorganization of function to the contralateral hemisphere – three out of four for words and three out of four for faces. By contrast, only two patients showed reorganization in the dorsal pathway – one patient for tools (patient FO) and one for global form (patient BI), regardless of whether the analyses were restricted to ROIs or the entire hemisphere. These findings are consistent with the only other known study that compared reorganization of ventral and dorsal functions, in a patient with resections to both pathways (Ahmad et al., 2022), and provides support for our initial hypothesis that the dorsal pathway may be less able to reorganize potentially because it matures earlier than the ventral pathway.

How do we reconcile these results with prior studies that have examined neural and behavioral recovery of perceptual functions following damage or resection? As described in the introduction, previous work has found evidence that the functions of the ventral pathway reorganize following VOTC resections (Liu et al., 2019; Liu et al., 2018), and that hemispherectomy patients retain a high degree of word and face recognition performance (Granovetter et al., 2022). However, it is important to note that not all patients in these studies showed evidence of reorganization (Liu et al., 2019). In fact, studies find mixed evidence of functional reorganization across the literature. For instance, many studies find no evidence that ventral functions, such as word and face recognition, recover following damage in childhood (Farah, Rabinowitz, Quinn, & Liu, 2000; Hadjikhani & de Gelder, 2002), even when the damage occurred on day 1 of life. By contrast, other studies find successful reorganization and normal recognition performance (Cohen et al., 2004; Mancini, de Schonen, Deruelle, & Massoulier, 1994), even when the disruptions occurred late in childhood (for review, see Liu & Behrmann, 2017; Vargha-Khadem & Polkey, 1992).

One important, but not universal, factor that seems to impact reorganization in previous studies is whether children experience unilateral or bilateral damage. On the whole, patients seem more likely to show neural reorganization and recovery of function if only one hemisphere was damaged, theoretically because the other hemisphere is able to compensate (Liu & Behrmann, 2017). Our results align with this literature on ventral pathway reorganization. We found that the majority, but not all, patients with hemispherectomy showed reorganization of function to their contralateral hemisphere in their ventral pathway. Given that only one hemisphere was resected, their intact hemisphere was available to compensate for the damage.

It is less clear, however, why patients showed abnormal responses to conditions that should already be lateralized to the intact hemisphere. For instance, only two patients with an intact left hemisphere showed normal word responses, and only one patient with an intact right hemisphere showed normal face responses. One possibility is that reorganization in the ventral pathway causes ‘neural crowding’ or competition between representations (Danguecan & Smith, 2019; Lidzba, Staudt, Wilke, & Kra geloh-Mann, 2006). For example, the presence of word representation in the right hemisphere encroaches on regions that would normally be exclusively face-selective, and vice versa for the presence of face representations in the left hemisphere. Indeed, prior work has shown that a patient with reorganization of words to their right hemisphere also had smaller than normal face ROIs (Liu et al., 2018), and over typical development, the emergence of VWFA in children correlates with an increasingly smaller face response in the left hemisphere (Behrmann & Plaut, 2020; Dehaene, 2005; Dundas, Plaut, & Behrmann, 2013; Nordt et al., 2021). However, it is important to note, that, although only a few patients showed normal selectivity for the preferred condition in their intact hemisphere, almost all patients (7 out of 8) showed ventral responses that were consistent with at least one hemisphere of controls.

There are far fewer studies examining the organization of the dorsal pathway following damage in childhood, with most focusing on processes related to visually guided action. Yet, action-related processes may be less likely to reorganize following damage given the strong one-to-one anatomical mapping between motor movements and the contralateral hemisphere (Granovetter et al., under revision; Schieber, 2001). Here we examined perceptual processes of the dorsal pathway, namely tool and global form representations, and found little evidence that these functions reorganize to their contralateral homologue. These findings are consistent with the hypothesis that the dorsal pathway matures early, and therefore has a smaller window of plasticity relative to the ventral pathway. Moreover, these findings align broadly with the ‘dorsal vulnerability hypothesis’ (Braddick et al., 2003), which posits that the dorsal pathway is particularly sensitive to disruption in childhood. Indeed, studies have shown widespread deficiencies in perceptual processes linked to the dorsal pathway in developmental disorders like CVI or William’s syndrome (Grinter et al., 2010; Macintyre-Be on et al., 2010). Children with these disorders rarely recover normal perceptual abilities. However, it is important to acknowledge that direct comparisons between our results and these disorders are challenging because, unlike disruptions from surgery or injury, these disorders also cause persistent bilateral, brain-wide effects that may not be solely linked to the dorsal pathway.

It is interesting to note that the only two patients to show normal responses to global form in the dorsal pathway (patient BI and KT), are also the only patients who did not undergo a full anatomical hemispherectomy (see Table 1), and have some spared parietal tissue in their disrupted hemisphere. Although we did not observe any responses in their disrupted hemisphere (see Figure 6A), it is nevertheless possible that this spared tissue supports normal representations in their fully intact hemisphere. Indeed, BI is the only patient to show normal responses for both tools and global form in the dorsal pathway. Interestingly, she is also the only patient that does not show any normal responses in her ventral pathway. However, this pattern is less consistent for tools, such that only patient FO demonstrated evidence of reorganization in for tools, and, unfortunately, data on tools was not available for KT.

A final, and related, puzzle from our work is how hemispherectomy patients maintain relatively accurate behavioral performance despite some patients showing abnormal selectivity in ventral and dorsal pathways Indeed, all of the hemispherectomy patients tested are able to read and write (albeit, to a greater or lesser extent), and show good performance on word, face, and shape recognition tasks (see Methods and Supplemental Materials). One explanation is that some degree of neural response for each condition is sufficient, though not optimal, to support a base rate of behavioral performance. Indeed, although no patient showed normal selectivity across all the conditions tested, almost all patients showed at least some activation to each localizer, with many also showing above chance multivariate decoding – particularly in the dorsal pathway.

In conclusion, we sought to provide a detailed exploration of the capacity of ventral and dorsal pathways to reorganize following large-scale resections and to shed light on the possible developmental trajectories of each pathway. Using a within-subjects design, we found greater evidence of reorganization for the ventral pathway than the dorsal pathway, consistent with the claim that the ventral pathway matures later and has an extended window of plasticity, compared with the dorsal pathway. However, we also found evidence that hemispherectomy may disrupt preferred representations in patients’ intact hemisphere (perhaps as a result of neural crowding), which makes drawing overarching conclusions about the nature of neural reorganization in the visual system challenging. To successfully characterize the processes that drive functional reorganization, future research must use larger sample sizes and collect greater amounts of data per patient, so as to better relate patient-specific factors to neural processes.

## Acknowledgements

We thank the Pediatric Epilepsy Surgical Alliance for assistance in recruiting patients, especially Monika Jones, JD, Chief Executive Officer. The research was funded by a grant from the National Science Foundation grant (BCS2123069) to M.B, and from the National Institutes of Health (NEI R01EY027018) to M.B. and C. P. MB also acknowledges support from P30 CORE award EY08098 from the National Eye Institute, NIH, and unrestricted supporting funds from The Research to Prevent Blindness Inc, NY, and the Eye & Ear Foundation of Pittsburgh. S.R. is supported by the National Science Foundation Graduate Research Fellowship Program (DGE2140739). M.G. is supported by predoctoral fellowship award from American Epilepsy Society (#847556). V.A. is supported by the University of Pennsylvania MindCORE fellowship and the data-driven discovery (DDDI) fellowship.

---

**Supplemental Figure 1.**
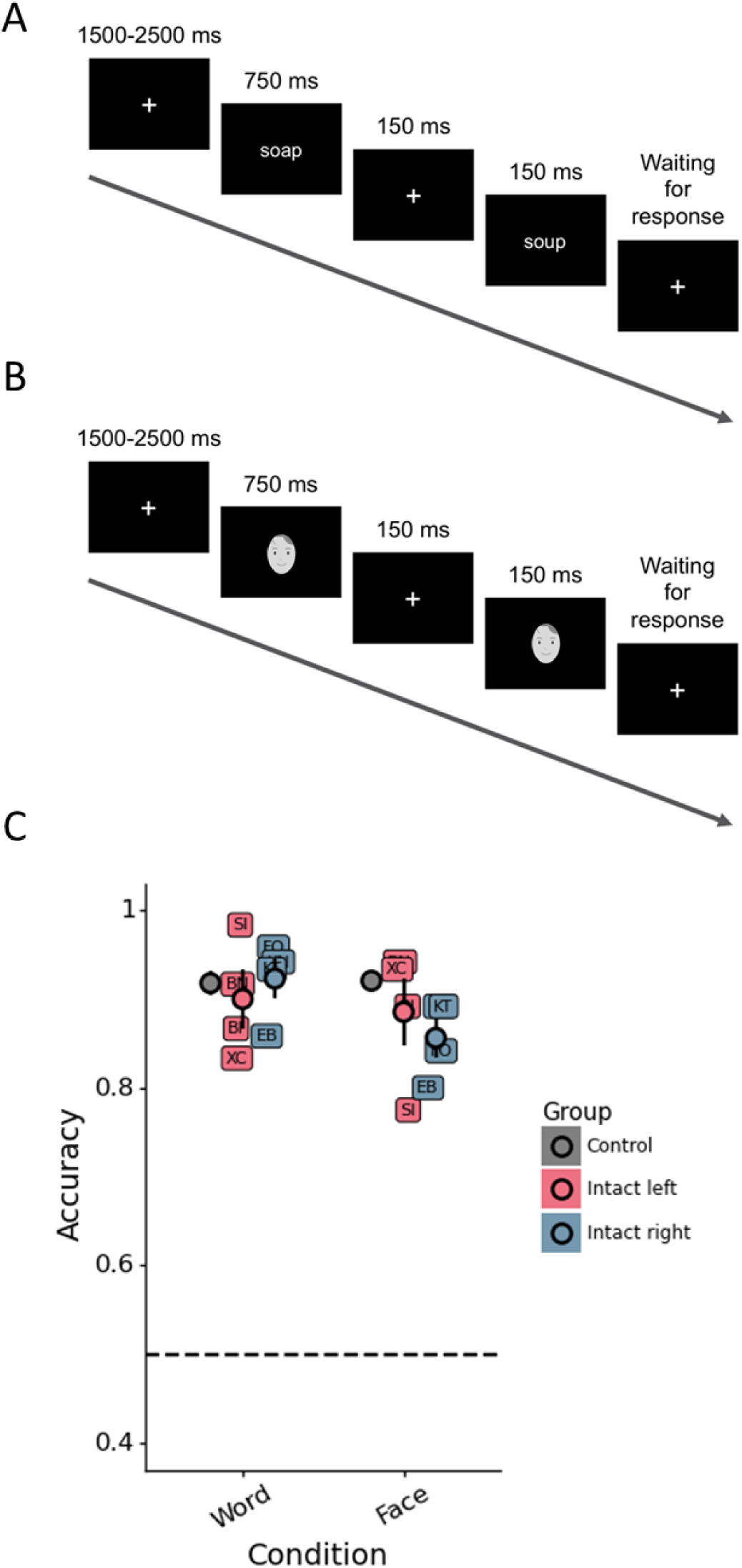
Performance on word and face discrimination task. (A-B) On each trial of the (A) word and (B) face discrimination tasks, participants were required to indicate whether sequentially presented words/faces were the same or different. (C) Word and face discrimination accuracy for controls, as well as patients with either intact left or right hemispheres. Note, in this example, face photographs were replaced with illustrations to comply with bioRxiv’s submission policy.

**Supplemental Figure 2.**
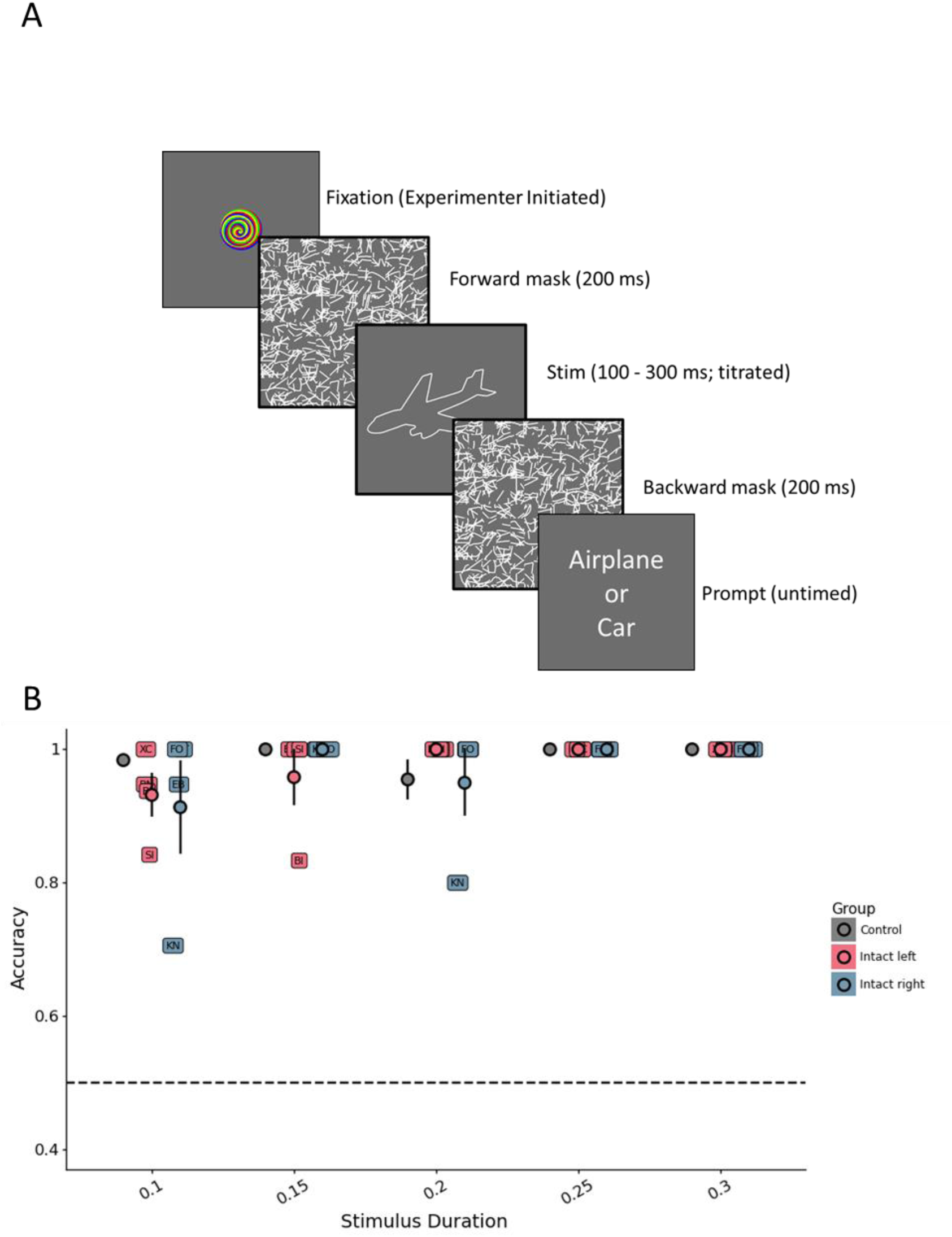
Performance on the shape recognition task. (A) On each trial of the object recognition task, participants were shown a rapidly presented shape (forward and backward masked) and then asked to report the identity of the object using a forced choice procedure. The distracting label in the prompt was a randomly selected object of the same animacy category. The stimulus presentation duration was titrated such that the duration decreased by 50 ms for every two trials participants answered correctly. (B) Shape recognition accuracy as a function of stimulus duration for controls and hemispherectomy patients with either an intact left or right hemisphere.

**Supplemental Figure 3.**
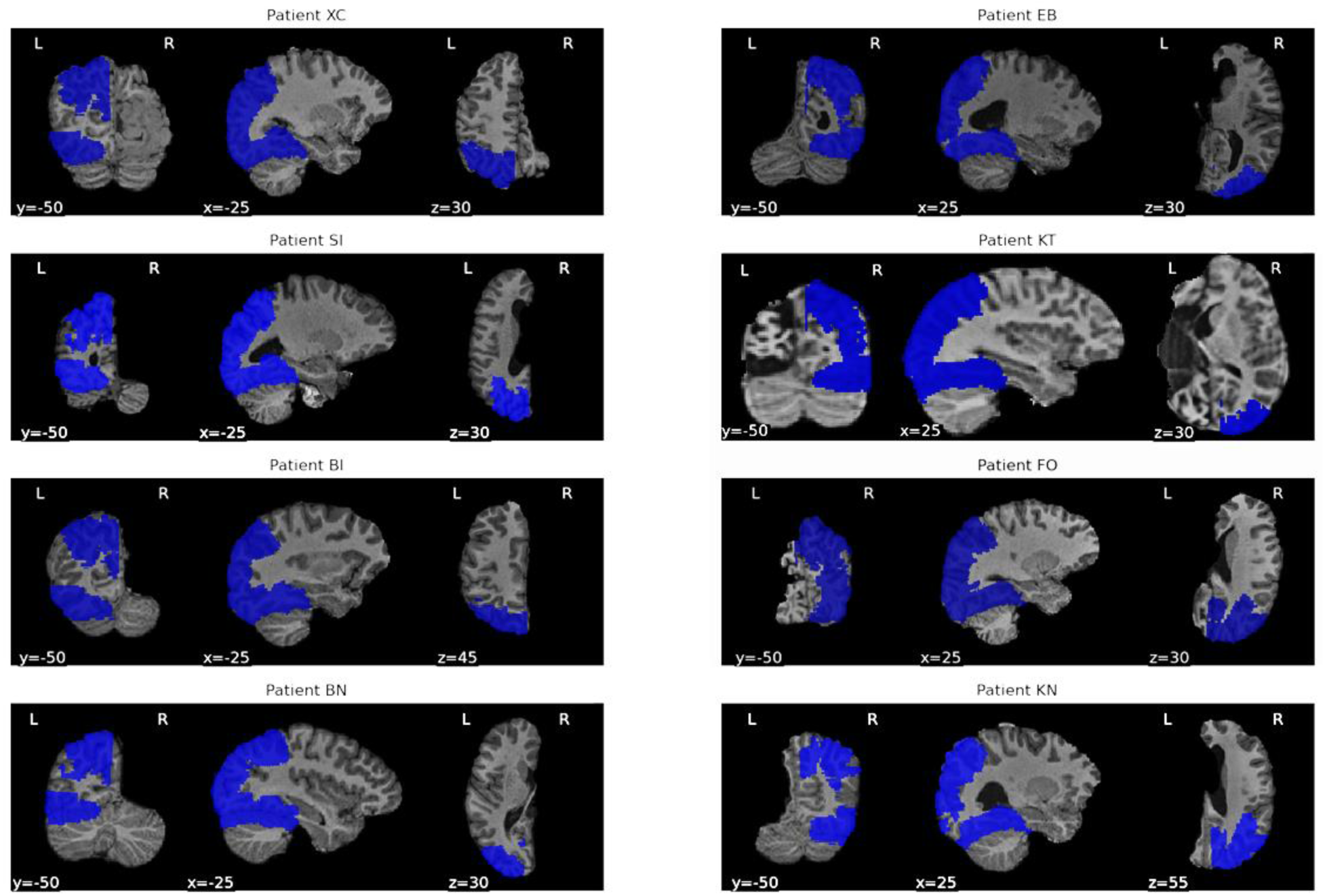
A visualization of the dorsal and ventral ROI masks overlaid on each patient’s individual anatomy.

**Supplemental Figure 4.**
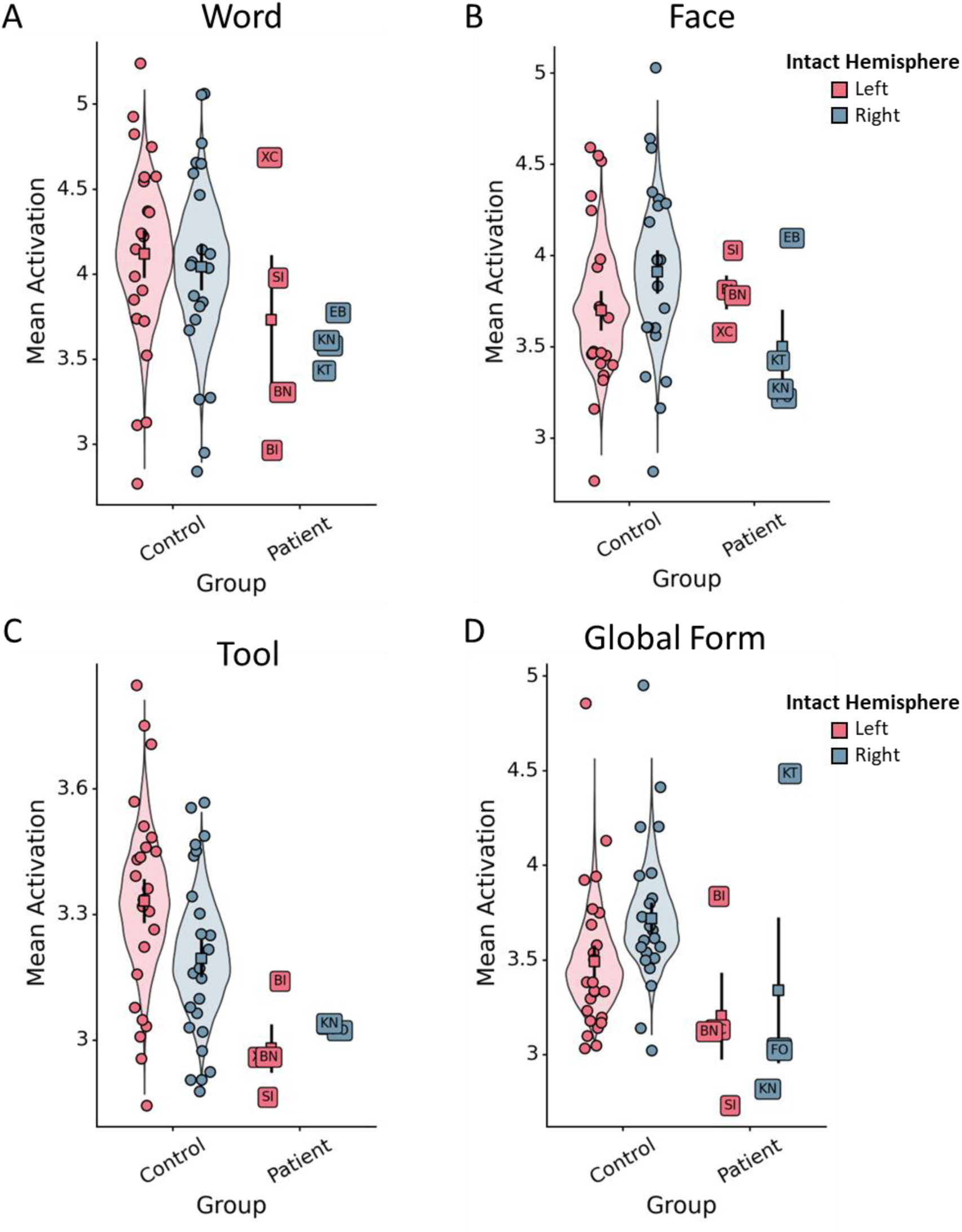
Mean activation for (A) words, (B) faces, (C) tools, and (D) global form in each patient and control’s hemisphere. Violin plots depict the bootstrapped distribution of control participants’ activation. Each unlabeled point refers to a single control participant’s data, and each labeled point corresponds to a single patient. Square points and error bar refer to the mean and standard error of each group, respectively.

**Supplemental Figure 5.**
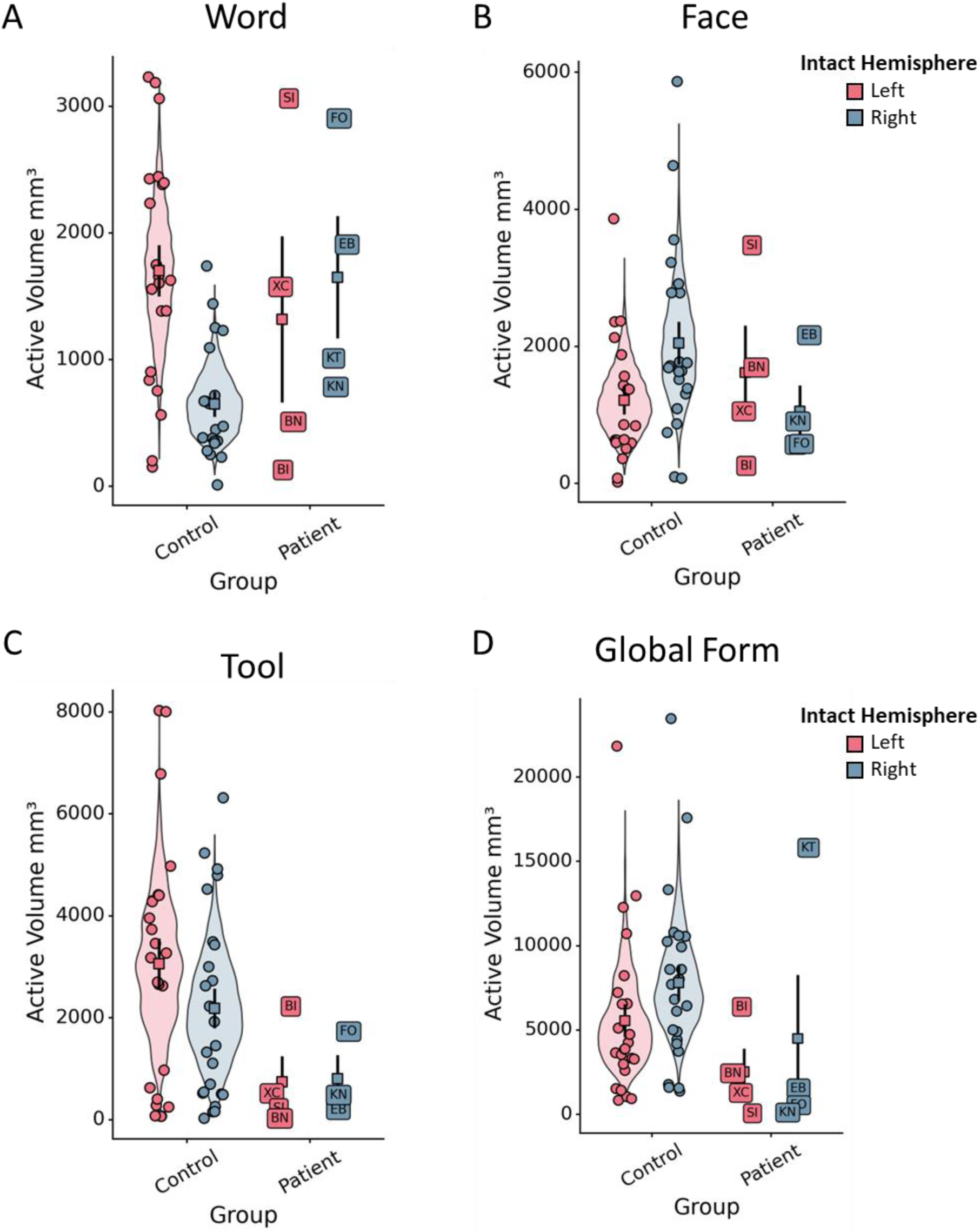
Mean active volume for (A) words, (B) faces, (C) tools, and (D) global form in each patient and control’s hemisphere. Violin plots depict the bootstrapped distribution of control participants’ active volume. Each unlabeled point refers to a single control participant’s data, and each labeled point corresponds to a single patient. Square points and error bar refer to the mean and standard error of each group, respectively.

**Supplemental Figure 6.**
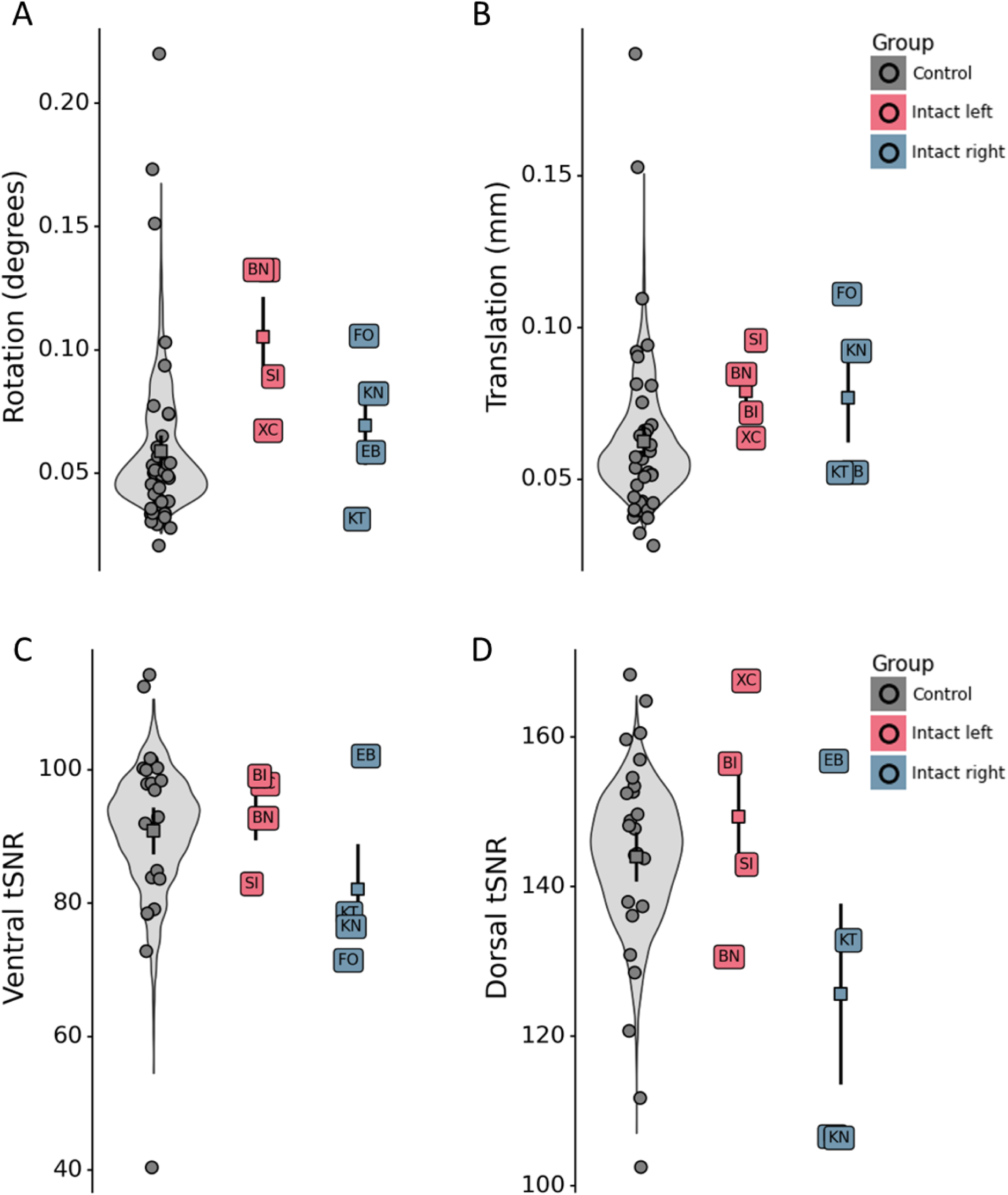
Signal quality metrics summarizing (A) rotation, (B) translation, (C) ventral temporal signal-to-noise ratio (tSNR), and (D) dorsal tSNR. Violin plots depict the bootstrapped distribution of control participants’ signal quality metrics. Each unlabeled point refers to a single control participant’s data, and each labeled point corresponds to a single patient. Square points and error bar refer to the mean and standard error of each group, respectively.

**Supplemental Table 1.** At-a-glance summary of selectivity results across the entire hemisphere. Each row indicates whether a patient’s summed selectivity score was below the control distribution (95% CIs). Asterisks (*) indicate that the patient’s score was below the control distribution, dashes (-) indicate that the patient’s score was inside or above the control distribution. ‘LH and ‘RH’ indicate whether the scores were compared to controls’ left hemisphere or right hemisphere, respectively. The ‘preferred’ hemisphere for each condition is underlined and in bold.

| Patient | Intact Hemisphere | Word |  | Face |  | Tool |  | Global Form |  |
| --- | --- | --- | --- | --- | --- | --- | --- | --- | --- |
|  |  | **<u>LH</u>** | RH | LH | **<u>RH</u>** | **<u>LH</u>** | RH | LH | **<u>RH</u>** |
| XC | Left | - | - | - | * | * | * | * | * |
| SI | Left | - | - | - | - | * | * | * | * |
| BI | Left | * | * | * | * | - | - | - | - |
| BN | Left | - | - | - | - | * | * | - | * |
| EB | Right | - | - | - | - | * | - | * | * |
| KT | Right | - | - | - | * | n/a | n/a | - | - |
| FO | Right | - | - | * | * | - | - | * | * |
| KN | Right | - | - | - | * | * | * | * | * |

**Supplemental Table 2.** At-a-glance summary of decoding results. Each row indicates whether a patient’s decoding accuracy was below the control distribution (95% CIs). Asterisks (*) indicate that the patient’s score was below the control distribution, dashes (-) indicate that the patient’s score was inside or above the control distribution. ‘LH and ‘RH’ indicate whether the scores were compared to controls’ left hemisphere or right hemisphere, respectively. The ‘preferred’ hemisphere for each condition is underlined and in bold.

| Patient | Intact Hemisphere | Word |  | Face |  | Tool |  | Global Form |  |
| --- | --- | --- | --- | --- | --- | --- | --- | --- | --- |
|  |  | **<u>LH</u>** | RH | LH | **<u>RH</u>** | **<u>LH</u>** | RH | LH | **<u>RH</u>** |
| XC | Left | - | - | - | - | - | - | - | - |
| SI | Left | - | - | - | - | * | - | * | - |
| BI | Left | * | * | * | * | * | * | - | - |
| BN | Left | * | * | * | - | - | - | * | * |
| EB | Right | - | - | - | - | - | - | - | - |
| KT | Right | - | - | * | * | n/a | n/a | - | - |
| FO | Right | - | - | * | * | * | * | * | * |
| KN | Right | * | * | * | * | - | - | - | - |

## Notes

### Competing Interest Statement

The authors have declared no competing interest.

